# *π*-PrimeNovo: An Accurate and Efficient Non-Autoregressive Deep Learning Model for De Novo Peptide Sequencing

**DOI:** 10.1101/2024.05.17.594647

**Authors:** Xiang Zhang, Tianze Ling, Zhi Jin, Sheng Xu, Zhiqiang Gao, Boyan Sun, Zijie Qiu, Nanqing Dong, Guangshuai Wang, Guibin Wang, Leyuan Li, Muhammad Abdul-Mageed, Laks V.S. Lakshmanan, Wanli Ouyang, Cheng Chang, Siqi Sun

## Abstract

Peptide sequencing via tandem mass spectrometry (MS/MS) is fundamental in proteomics data analysis, playing a pivotal role in unraveling the complex world of proteins within biological systems. In contrast to conventional database searching methods, deep learning models excel in de novo sequencing peptides absent from existing databases, thereby facilitating the identification and analysis of novel peptide sequences. Current deep learning models for peptide sequencing predominantly use an autoregressive generation approach, where early errors can cascade, largely affecting overall sequence accuracy. And the usage of sequential decoding algorithms such as beam search suffers from the low inference speed. To address this, we introduce ***π***-PrimeNovo, a non-autoregressive Transformer-based deep learning model designed to perform accurate and efficient de novo peptide sequencing. With the proposed novel architecture, ***π***-PrimeNovo achieves significantly higher accuracy and up to 69x faster sequencing compared to the state-of-the-art methods. This remarkable speed makes it highly suitable for computation-extensive peptide sequencing tasks such as metaproteomic research, where ***π***-PrimeNovo efficiently identifies the microbial species-specific peptides. Moreover, ***π***-PrimeNovo has been demonstrated to have a powerful capability in accurately mining phosphopeptides in a non-enriched phosphoproteomic dataset, showing an alternative solution to detect low-abundance post-translational modifications (PTMs). We suggest that this work not only advances the development of peptide sequencing techniques but also introduces a transformative computational model with wide-range implications for biological research.

## Introduction

Protein identification plays a critical role in proteomics, and currently, shotgun proteomics via mass spectrometry is widely acknowledged as the primary method for this purpose [1]. This technique involves enzymatically digesting proteins into peptides, which are subsequently subjected to mass spectrometry analysis, resulting in tandem mass spectra. These spectra are invaluable for gaining insights into the sequences and structures of the peptides. A fundamental aspect of protein identification involves deducing the amino acid sequence from the tandem mass spectra [2]. At present, database searching is the primary approach for peptide and protein identification, utilizing tools such as SEQUEST [3], Mascot [4], MaxQuant/Andromeda [2], PEAKS DB [5], and pFind [6]. However, the effectiveness of these methods relies on the availability of a comprehensive sequence database that encompasses all potential proteins in the sample. This requirement limits their applicability in scenarios such as monoclonal antibody sequence assembly [7], the identification of novel antigens [8], and the sequencing of metaproteomes lacking established databases [9].

Over the last two decades, a wide array of de novo sequencing tools for peptides have emerged and contributed to the field’s advancement [8, 10–21]. The core idea behind these algorithms is to deduce amino acid compositions and modifications by analyzing the mass differences between fragment ions. Initial de novo sequencing algorithms applied graph theory and dynamic programming. For instance, PepNovo [11] took spectra as graphs, with nodes representing fragment ions connected based on mass differences equal to an amino acid’s mass. PEAKS [10] employed a dynamic programming approach, awarding or penalizing the choice of amino acids based on the presence or absence of fragment ions. DeepNovo [12] marked a significant evolution by introducing deep learning into the realm of de novo peptide sequencing. It combined a convolutional neural network (CNN) to extract data from spectrum peaks with a long shortterm memory network (LSTM) for processing peptide sequences. To better handle high-precision mass spectrometry data, PointNovo [13] introduced a model with an order-invariant network to learn the features in a spectrum. Following the success of Transformers in language translation [22], Casanovo [15] applied a similar approach to peptide sequencing, treating it as a translation task from spectrum peak sequences to amino acid sequences. The authors of Casanovo also retrained Casanovo V2 [16] on a large dataset containing 30 million labeled spectra. Most recently, PepNet [19] employed a fully convolutional neural network for fast de novo peptide sequencing. To tackle the missing-fragmentation problem, where the given spectrum does not contain most of the theoretical peaks of the true peptide, GraphNovo [23] introduced a two-stage de novo peptide sequencing algorithm based on a graph neural network. Despite these advancements [17–21, 23], the peptide recall of deep learning-based de novo sequencing algorithms in shotgun proteomics, defined as the percentage of correctly predicted spectra among all spectra, remains far from optimal, ranging from 30% to 50% on standard nine-species benchmark.

Currently, all deep learning models for de novo peptide sequencing are based on the autoregressive framework [24], meaning the generation of each amino acid is heavily reliant on its predicted predecessors, resulting in a unidirectional generation process. However, the significance of bidirectional information is paramount in peptide sequencing, as the presence of an amino acid is intrinsically linked to its neighbors in both directions [21]. In autoregressive models, any errors in early amino acid predictions can cascade, affecting subsequent generations. Autoregressive decoding algorithms such as beam search lack the capability to retrospectively modify previously generated content, making it challenging to control the total mass of the generated sequence. This limitation arises because each token is produced based on its predecessor, meaning that altering any previously generated token would consequently shift the distribution of subsequent tokens and therefore require a re-generation of the whole sequence [25].

In this research, we introduce *π*-PrimeNovo (shortened as PrimeNovo), representing a significant departure from conventional autoregressive approaches by adopting a non-autoregressive approach to effectively address the unidirectional problems of autoregressive methods. This innovation stands as the first non-autoregressive Transformer-based model in this field. Such design enables a simultaneous sequence prediction, granting each amino acid a comprehensive bidirectional context. Another key advancement in PrimeNovo is the integration of a novel Precise Mass Control (PMC) unit, uniquely compatible with the non-autoregressive framework, which utilizes precursor mass information to generate controlled and precise peptide sequences. This precise mass control, coupled with bidirectional generation, significantly enhances peptide-level performance.

PrimeNovo consistently demonstrates impressive peptide-level accuracy, achieving an average peptide recall of 64% on the widely used nine-species benchmark dataset. This performance significantly surpasses the existing best model, which achieves a peptide recall of 54% [16]. Across a diverse range of other MS/MS datasets, PrimeNovo consistently maintains a notable advantage in peptide recall over the state-of-the-art model, achieving relative improvements from 16% to even doubling the accuracy, which highlights its exceptional performance and reliability. Moreover, by avoiding the sequential, one-by-one generation process inherent in autoregressive models, PrimeNovo also substantially increases its inference speed. This acceleration is further enhanced through the use of dynamic programming and CUDA-accelerated computation, allowing PrimeNovo to surpass the existing autoregressive models by up to 69 times. This speedup advantage enables PrimeNovo to make accurate predictions on large-scale spectrum data. We have demonstrated that PrimeNovo excels in large-scale metaproteomic research by accurately identifying a significantly greater number of species-specific peptides compared to previous methods, reducing the processing time from months, as required by Casanovo V2 with beam search, to just days. Furthermore, PrimeNovo’s versatility extends to the identification of PTMs, showcasing its potential as a transformative tool in proteomics research.

## Results

### PrimeNovo sets new benchmark with 64% peptide recall, achieving over 10% improvement in widely-used nine-species dataset

Echoing the approach of Casanovo V2, we utilized the large-scale MassIVE-KB dataset [26], featuring around 30 million Peptide-to-Spectrum Matches (PSMs), as our training data. PrimeNovo was then evaluated on the nine-species testing benchmark directly. It is crucial to note, however, that baseline models like PointNovo, DeepNovo, and Casanovo were originally trained using the leave-one-species-out cross-validation (CV) strategy [27] on the nine-species dataset. This strategy involves training on eight species and evaluating on the ninth each time. To facilitate a fair comparison, we also trained PrimeNovo on the nine-species dataset using the same CV strategy, following the data split used by all other baseline models. As shown in Fig. 2a, PrimeNovo CV outperformed other baseline models trained with this strategy by a large margin. Notably, even when trained solely on the nine-species benchmark dataset, PrimeNovo CV already matched the performance of Casanovo V2, which is the model trained on the large-scale MassIVE-KB dataset. When trained on the MassIVE-KB dataset, PrimeNovo set new state-of-the-art results across all species in the nine-species benchmark (Fig. 2b and Supplementary Fig. 4). The average peptide recall improved significantly, increasing from 45% with Casanovo to 54% with Casanovo V2, and further to 64% with PrimeNovo. This marks a 10% improvement over Casanovo V2 and a 19% increase over Casanovo. In the recall-coverage curve (Fig. 2a), PrimeNovo consistently held the top position across all coverage levels and species, reaffirming its status as a leading model in de novo peptide sequencing. Additionally, we tested PrimeNovo on a revised nine-species test set introduced by Yilmaz [16], which featured higher data quality and a larger quantity of spectra, covering a wider range of data distributions for each species. In this updated test, PrimeNovo’s average peptide recall soared to 75% across all species, from the previous 65% by Casanovo V2. A detailed comparison of these results is available in Supplementary Fig. 2. The outcomes from both the original and revised nine-species benchmark datasets highlight PrimeNovo’s capability to accurately predict peptides across various species, demonstrating its effectiveness and versatility.

**Fig. 1.**
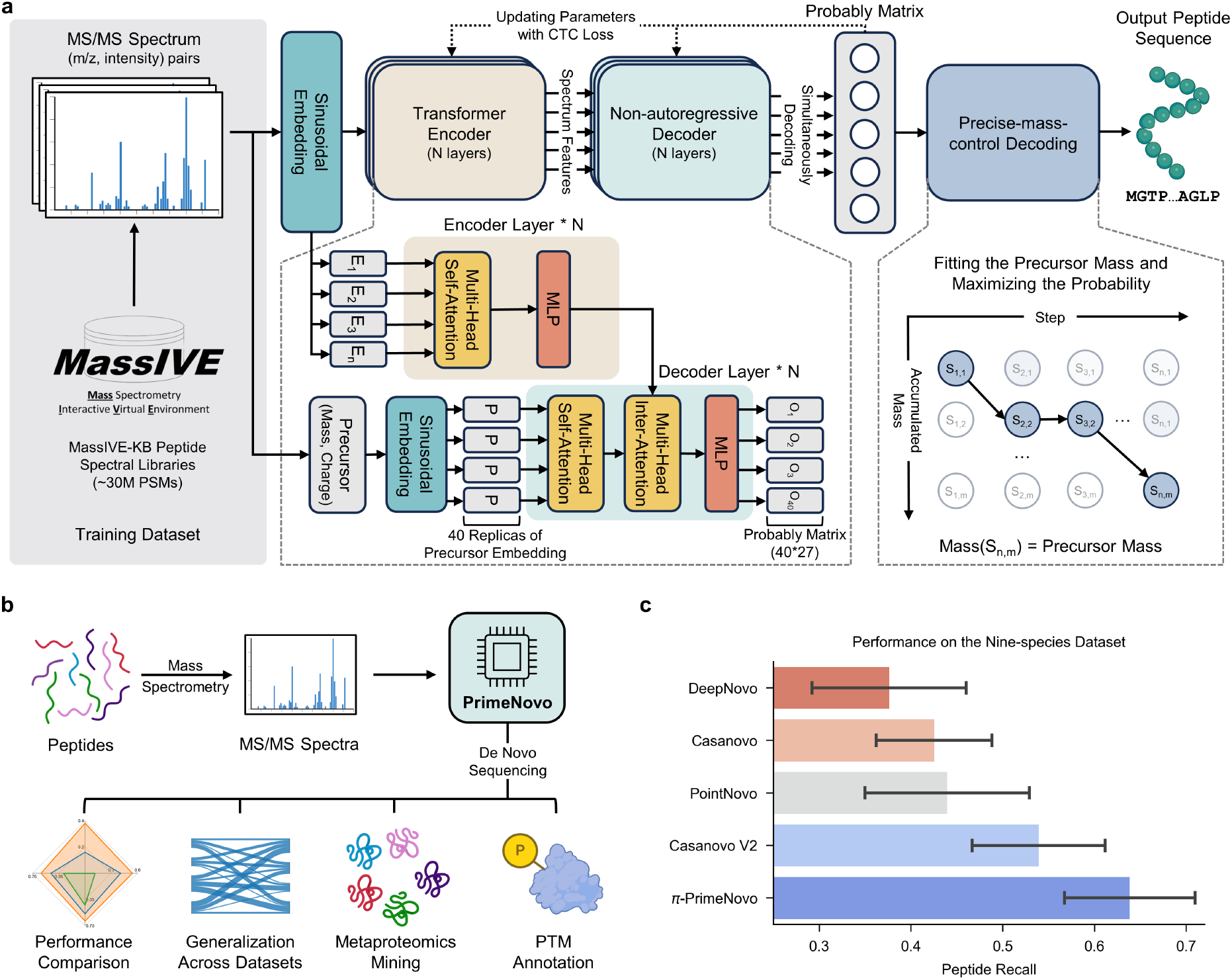
PrimeNovo stands as the pioneering biological non-autoregressive Transformer model, delivering precise peptide sequencing. **a** Model Architecture Overview: Our model takes MS/MS spectra as input and generates the predicted peptide sequence. It comprises two key components: 1) a non-autoregressive Transformer model backbone optimized with Connectionist Temporal Classification (CTC) loss, enabling simultaneous amino acid prediction at all positions. 2) The Precise Mass Control (PMC) decoding unit, which utilizes predicted probabilities to precisely optimize peptide generation to meet mass requirements. **b**. Applications and Biological Insights: PrimeNovo’s capabilities extend to downstream tasks and offer valuable insights for various biological investigations. **c**. Average Performance Comparison: This chart illustrates the average performance of PrimeNovo alongside four other top-performing models on the widely utilized nine-species benchmark dataset. Each bar represents the mean peptide recall for the respective approach. The black line indicates the 95% confidence interval. Notably, results for DeepNovo, Casanovo, and Casanovo V2 are based on model weights released by the original authors, while PointNovo’s results are cited from the published work, as the original model weights were not shared by PointNovo’s authors.

**Fig. 2.**
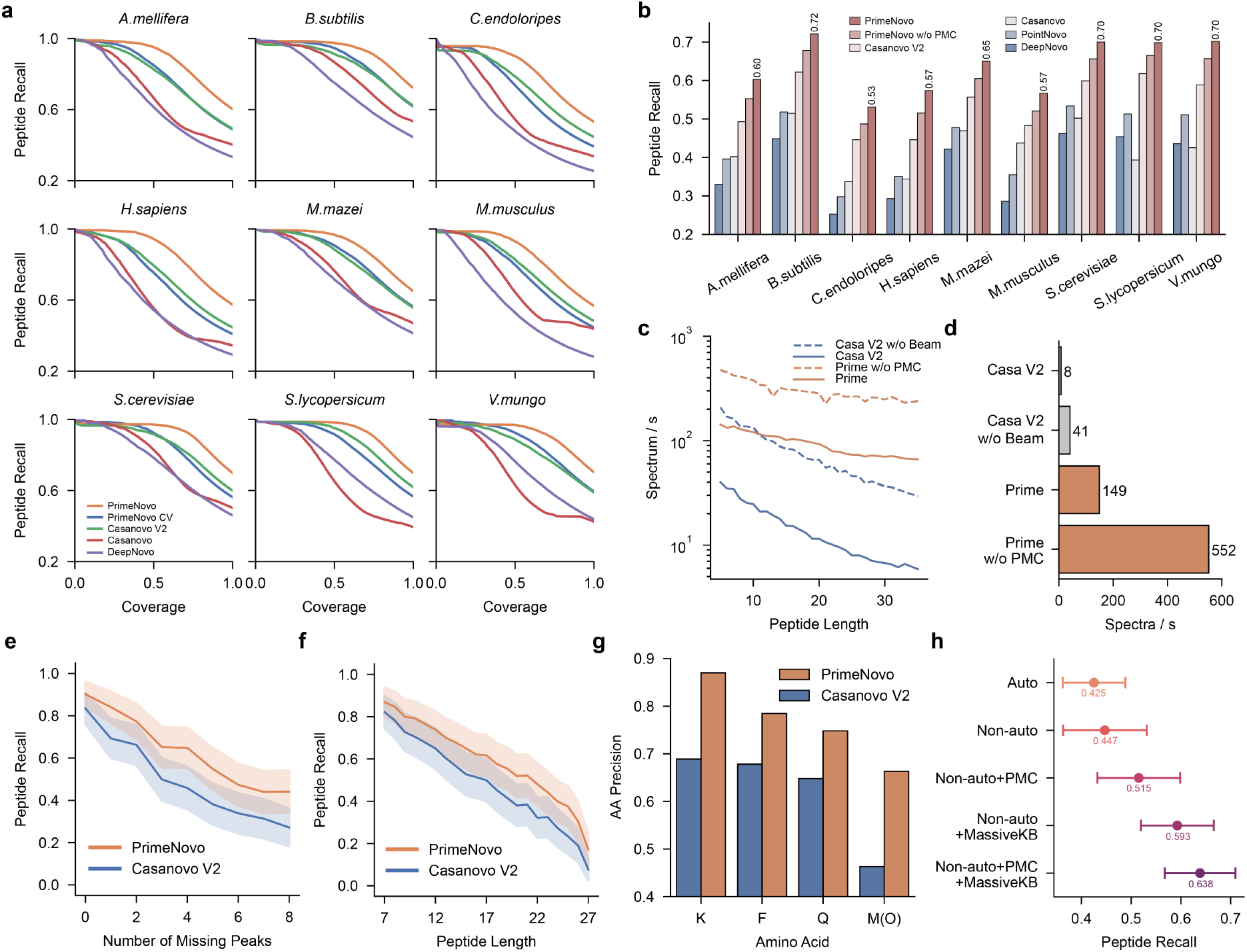
A detailed comparison between PrimeNovo and previous deep learning-based approaches on the nine-species benchmark dataset. **a**. The performance comparison between PrimeNovo and other de novo algorithms for recall-coverage curves on the nine-species benchmark dataset. These curves illustrate recall (the averaged peptide recall) – coverage (the proportion of the predicted spectra to all spectra ranked by the model’s confidence) relationships across all confidence levels for each test species. PrimeNovo CV represents our model trained on the nine-species benchmark dataset using a cross-validation strategy. PrimeNovo represents our model trained on the MassIVE-KB dataset. **b**. The average prediction performance on each individual species for PrimeNovo and comparison models. PrimeNovo w/o PMC presents results obtained using CTC beam search decoding without PMC. **c**. Inference speed with respect to the prediction length between Casanovo V2 and PrimeNovo, with and without a post-decoding strategy for each model. **d**. Inference Speed Comparison: A comparison of inference speeds, measured in the number of spectra decoded per second, between PrimeNovo and Casanovo V2. The speed tests were conducted on the same computational hardware (single A100 NVIDIA GPU) and averaged over data from all test species. **e-f**. Influence of Missing Peaks and Peptide Length: These plots reveal how the degree of missing peaks and the length of true labels affect the predictions of PrimeNovo and Casanovo V2. **g**. Performance on Amino Acids with Similar Masses: A comparison of Casanovo V2 and PrimeNovo in predicting amino acids with very similar molecular masses, such as K (128.094963) with Q (128.058578) and F (147.068414) with Oxidized M (147.035400). **h**. Ablation Study: An analysis of the impact of adding each module of our approach on the overall performance.

PrimeNovo, leveraging its bi-directional information integration and parallel generation process as a non-autoregressive model, convincingly establishes its superiority across various facets of sequencing tasks, transcending mere high prediction accuracy. Firstly, our non-autoregressive model offers a substantial improvement in the inference speed compared to the autoregressive models of similar sizes, thanks to its concurrent generation process. As depicted in Fig. 2d, PrimeNovo, even without the Precise Mass Control (PMC) unit, achieves a staggering speed advantage of 13.5 times faster over Casanovo V2 without beam search decoding under identical testing conditions (*i*.*e*., using the same machine with identical CPU and GPU specifications). Upon incorporating post-prediction decoding strategies (PMC for PrimeNovo and beam search for Casanovo V2), PrimeNovo’s advantage in inference speed becomes even more pronounced, making it over 18 times faster than Casanovo V2. Notably, considering that PrimeNovo without PMC can already outperform Casanovo V2 with beam search by an average of 6% on the nine-species benchmark dataset (as demonstrated in Fig. 2b), users can experience a maximum speedup of 69 times while making only minimal sacrifices in prediction accuracy when PMC is not deployed. It is also noteworthy that the generation speed of autoregressive models is significantly affected by the length of the generated sequence, unlike our model. Fig. 2c illustrates that PrimeNovo exhibits negligible speed degradation, which is partly due to encoding the longer spectrum input for longer sequence, as the peptide length increases, whereas Casanovo V2 experiences a quadratic decrease in speed as sequences grow longer.

Furthermore, PrimeNovo exhibits exceptional prediction robustness across various challenges, including different levels of missing peaks in the spectrum, varying peptide lengths, and amino acid combinations that are prone to confusion. To illustrate this robustness, we categorized predictions on the nine-species benchmark dataset based on the degree of missing peaks in the input spectrum and the number of amino acids in the target peptide. The calculation of missing peaks in each spectrum follows the methodology outlined in a previous study by [7], where we compute all the theoretical *m/*z values for potential *y* ions and *b* ions based on the true label and determine how many of these theoretical peaks are absent in the actual spectrum. As presented in Fig. 2e, it’s not surprising to observe a decline in prediction accuracy as the number of missing peaks in the spectra increases. However, PrimeNovo consistently indicates superior performance across all levels of missing peaks and consistently outperforms Casanovo V2. Similarly, Fig. 2f illustrates that PrimeNovo maintains its higher accuracy compared to Casanovo V2, irrespective of the length of the peptide being predicted. In Fig. 2g, we further observe that PrimeNovo excels in accurately predicting amino acids that are challenging to identify due to their closely similar mass (less than 0.05 Da) to other amino acids. Specifically, the aa precision of all four similar amino acids is more than 10% more accurate on average compared to that of Casanovo V2. Specifically, the precision advantage is more than 18% on both K and Oxidized M amino acids.

We then conducted an ablation study to investigate the performance gains achieved by each component of our model on the nine-species benchmark dataset. From Fig. 2h, we observe 2% improvement in peptide recall when transitioning from an autoregressive model to a non-autoregressive model. The gain in performance is magnified by a large amount (7%) when PMC is introduced, as controllable generation is important in such tasks and improves the accuracy of our generated sequence. Remarkably, the performance boost from the non-autoregressive model is most pronounced when transitioning from the CV training data to the MassIVE-KB dataset, as the substantial increase in training data proves invaluable for learning the underlying bi-directional patterns in the sequencing task. Lastly, we see that utilizing PMC with augmented training data achieves the highest prediction accuracy, which further demonstrates PMC’s importance under different data availability situations.

### PrimeNovo exhibits strong generalization and adaptability capability across a wide array of MS/MS data sources

As MS/MS data can vary significantly due to differences in biological samples, mass spectrometer parameters, and post-processing procedures, there is often a substantial degree of distributional shift across various MS/MS datasets. To demonstrate PrimeNovo’s ability to generalize effectively across a wide spectrum of distinct MS/MS data for diverse downstream tasks, we conducted an evaluation of PrimeNovo’s performance on some of the most widely used publicly available MS/MS datasets. We then compared the results with those of the current state-of-the-art models, Casanovo and Casanovo V2. In addition to the nine-species benchmark dataset discussed earlier, we selected three prominent MS/MS datasets that represent varying data sources and application settings: the PT [28], IgG1-Human-HC [27], and HCC [29] datasets^1^.

We start by evaluating PrimeNovo’s ability to perform well in a “zero-shot” scenario, which means the model is tested without any specific adjustments to match the characteristics and distribution of the target dataset. As depicted in Fig. 3a and Supplementary Fig. 6, PrimeNovo exhibits significant performance superiority over both Casanovo V2 and Casanovo in terms of peptide recall when directly tested on three distinct datasets. Specifically, PrimeNovo outperforms Casanovo V2 by 13%, 14%, and 22% on PT, IgG1-Human-HC, and HCC datasets, respectively. This performance gap widens to 30%, 43%, and 38% when compared to Casanovo. For the IgG1-Human-HC dataset, following [7], we present the evaluation results for each human antigen type, as illustrated in Fig. 3b. PrimeNovo consistently outperforms Casanovo V2 across all six antigen types, achieving increased peptide recall ranging from 9% to 20%. We further examine the amino acid level accuracy on the unseen dataset. From Fig. 3c, it’s notable that PrimeNovo has a dominant AA level precision advantage over Casanovo V2 across all confidence levels of the model output. This indicates PrimeNovo’s better prediction of amino acids’ presence and locations.

**Fig. 3.**
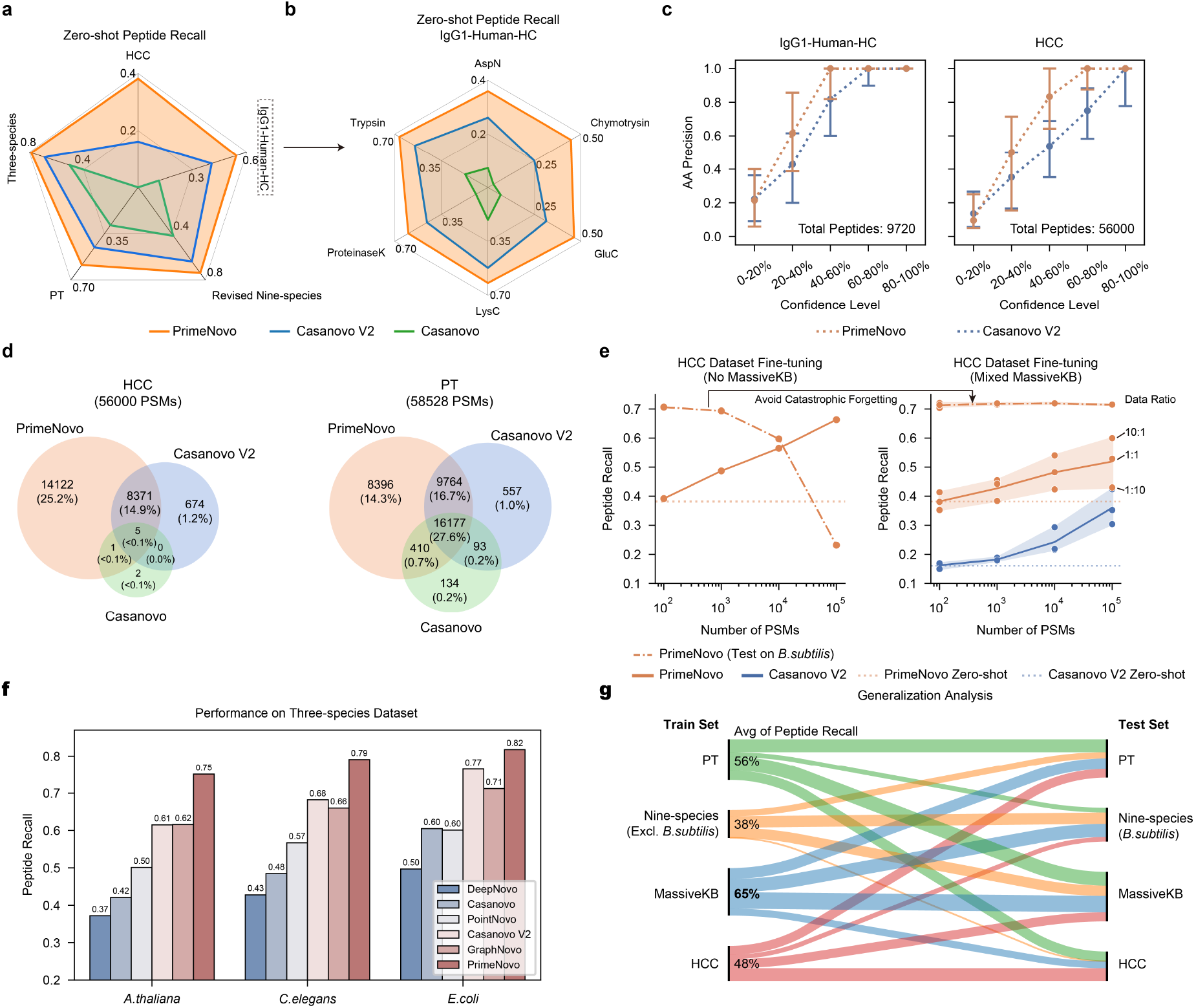
PrimeNovo’s exceptional performance extends to unseen spectra from various biological sample sources. **a**. Average Peptide Recall: This section details the average peptide recall of PrimeNovo compared to baseline models across four distinct large-scale MS/MS datasets. **b**. Enzyme-Specific Performance: Performance breakdown among six different proteolytic enzymes in the IgG1-Human-HC dataset. **c**. Amino Acid-Level Precision: The chart depicts the amino acid-level precision for PrimeNovo and Casanovo V2 on the IgG1-Human-HC and HCC datasets. The x-axis shows the coverage rate of predicted peptides based on each model’s confidence score. **d**. A Venn diagram illustrates the number of overlapping peptides among three de novo sequencing models and a traditional database searching algorithm. Each count represents identical peptides identified by both MaxQuant and the respective model for the same spectrum. **e**. Model Fine-Tuning Results: This chart demonstrates how performance on the HCC test dataset changes with the addition of more HCC training data during fine-tuning. The left side shows fine-tuning with only the HCC dataset, leading to catastrophic forgetting of the original data distribution (*nine-species benchmark dataset*). The right side shows fine-tuning with a mix of HCC and MassIVE-KB training data. **f**. A comparison of performance between PrimeNovo and five other de novo models on a 3-species test dataset. **g**. This diagram evaluates the model’s generalization capability when trained exclusively with each training dataset. The numbers indicate the average peptide recall when tested on all other datasets, highlighting the distributional transferability of each. The model trained on MassIVE-KB exhibited the highest average peptide recall, 65% (bolded).

To further assess the performance disparities under the zero-shot setting, we leveraged identified PSMs from MaxQuant in each dataset as the benchmark. Then we compared the number of overlapping PSMs between the predicted PSMs generated by each de novo algorithm and the PSMs identified by MaxQuant. As displayed in Fig. 3d, Casanovo performed poorly on the HCC dataset, with only 8 PSMs overlapping with MaxQuant. In contrast, Casanovo V2 identified 9050 overlapping PSMs, while PrimeNovo predicted up to 22499 PSMs that perfectly matched those identified by MaxQuant. On the PT dataset, PrimeNovo, Casanovo V2, and Casanovo had 34747, 26591, and 16814 overlapping PSMs with MaxQuant search results, respectively. PrimeNovo demonstrates a much more consistent prediction behavior, aligning closely with high-quality traditional database-searching peptide identification software.

Next, we examine how well PrimeNovo generalizes under the fine-tuning setting, which involves quickly adapting the model to new training data from the target distribution without starting the training process from scratch. This approach allows the model to leverage its previously acquired knowledge from the large dataset it was originally trained on and apply it to a more specific task or domain with only a minimal amount of additional training. We fine-tuned PrimeNovo on both the PT and HCC training datasets to assess the model’s adaptability. In order to gauge the impact of the quantity of additional data on finetuning performance, we conducted the fine-tuning with 100, 1000, 10000, and 100000 additional data points, respectively. We also fine-tuned Casanovo V2 under identical settings to compare the adaptability of the two models fairly. As depicted on the right side of Fig. 3e, augmenting the amount of additional data for fine-tuning does indeed enhance the model’s prediction accuracy on the corresponding test set, as the model gains a better understanding of the distributional nuances within the data. In comparison, PrimeNovo demonstrates a more robust ability to adapt to new data distributions and achieves higher accuracy after fine-tuning compared to the zero-shot scenario. It consistently outperforms Casanovo V2 when subjected to the same fine-tuning conditions, with 18% and 12% higher peptide level recall on HCC and PT test sets respectively when the fine-tuning reaches the best performance (Fig. 3e). It is noteworthy that a noticeable improvement in prediction accuracy is only observed after incorporating 10,000 additional MS data points during the fine-tuning process, indicating a recommended data size for future fine-tuning endeavors involving other data distributions.

It’s important to note that the fine-tuning process can lead the model to forget the original data distribution from the training set, which is referred to as catastrophic forgetting. As illustrated in the left part of Fig. 3e, when fine-tuning is conducted exclusively with the target data, the performance in the nine-species benchmark dataset experiences a significant and gradual decline as more data samples are included (indicated by the dashed line). However, when the target data is mixed with the original training data, catastrophic forgetting is mitigated, as evident from the dashed line in the right part of Fig. 3e. Indeed, fine-tuning exclusively with the target data does introduce a relatively higher performance gain in the target test set compared to fine-tuning with mixed data (solid line in Fig. 3e), where the difference can be as much as 15% when the amount of the new data used for fine-tuning is large.

By fine-tuning the model using a single dataset and then testing it on others, we can explore the similarities and disparities in data distributions among different pairs of datasets. This approach provides valuable insights into how closely related each MS/MS dataset is to the others and the extent to which a model’s knowledge can be transferred when trained on one dataset. In Fig. 3g, it’s not surprising to observe that the model exhibits the strongest transferability when the training and testing data share the same data source. Notably, MassIVE-KB, the training set for both our model and Casanovo V2, demonstrates the highest average peptide recall of 65% across all other test sets. This can be attributed to the diverse range of MS/MS data sources encompassed within the MassIVE-KB dataset, covering a wide spectrum of distinct MS/MS data. The PT dataset, with an average peptide recall of 56%, is also considered a high-quality dataset with robust transferability. It has been employed in the training of numerous other de novo models [21]. However, the models trained on the HCC and nine-species benchmark datasets do not generalize well to other testing datasets. The nine-species benchmark exclusively covers MS/MS data for the included nine species and has a relatively small data size, while the HCC dataset is specific to human hepatocellular carcinoma. Additionally, we observe that models trained with the nine-species benchmark dataset and MassIVE-KB datasets exhibit relatively poor performance when applied to the HCC dataset, suggesting a notable disparity in their data distribution.

Finally, we conduct a comparative analysis between PrimeNovo and concurrent approaches in de novo sequencing to illustrate the advancements and effectiveness of our method. Our comparative models, namely GraphNovo, a graph-based neural network, and PepNet, a CNN-based neural network, approach the problem from distinct angles, utilizing the latest deep learning techniques. It’s worth noting that both GraphNovo [23] and PepNet [19] are trained on their own designated training and testing datasets for their respective model versions. Consequently, we adopt a zero-shot evaluation approach, testing PrimeNovo on each of their test sets and comparing the results with their reported performances. We carefully examined the used data and ensured that there was no overlap between our training dataset and the test sets used by GraphNovo and PepNet. For the 3-species test set employed by GraphNovo, PrimeNovo demonstrates remarkable improvements in peptide recall, surpassing GraphNovo by 13%, 13%, and 11% in the A. thaliana, C. elegans, and E. coli species, respectively (see Fig. 3f). Furthermore, when tested on the PepNet test set, PrimeNovo exhibits a notable advantage of 14% and 24% in peptide recall over PepNet when predicting the peptide with charges of 2 and 3 respectively, detailed results of which are in Supplementary Fig. 11.

### PrimeNovo’s behavior analysis reveals an effective error correction mechanism behind non-autoregressive modeling and PMC unit

To gain a comprehensive understanding of the model’s behavior and to analyze how PrimeNovo utilizes the spectrum data to arrive at its final results, we employ some of the most recent model interpretability techniques, examining each component of our model in detail. We commence by visualizing the attention behavior of the encoder network in PrimeNovo and comparing it to that of Casanovo V2. The encoder’s role is critical, as it is responsible for feature extraction from the spectrum, significantly influencing how well the model utilizes input spectrum data. As depicted in the attention map in Fig. 4a, it is evident that Casanovo V2 predominantly assigns most of its attention weights to the first input position (the special token added at the beginning of the peak tokens). Attention weights for the remaining tokens are sparse, insignificant, and primarily concentrated along the diagonal direction. This behavior suggests that Casanovo V2 encodes information primarily within its special token, with limited utilization of other peak positions. In contrast, PrimeNovo exhibits a well-distributed attention pattern across different input peaks, each with varying levels of information density. Furthermore, we observe that the attention of PrimeNovo is more heavily allocated to peaks corresponding to the b-y ions of the true label, which are among the most crucial pieces of information for decoding the spectrum (as detailed in Supplementary Fig. 15. This highlights PrimeNovo’s capacity to extract information more effectively from tokens it deems essential, and this behavior remains consistently active across all nine layers.

**Fig. 4.**
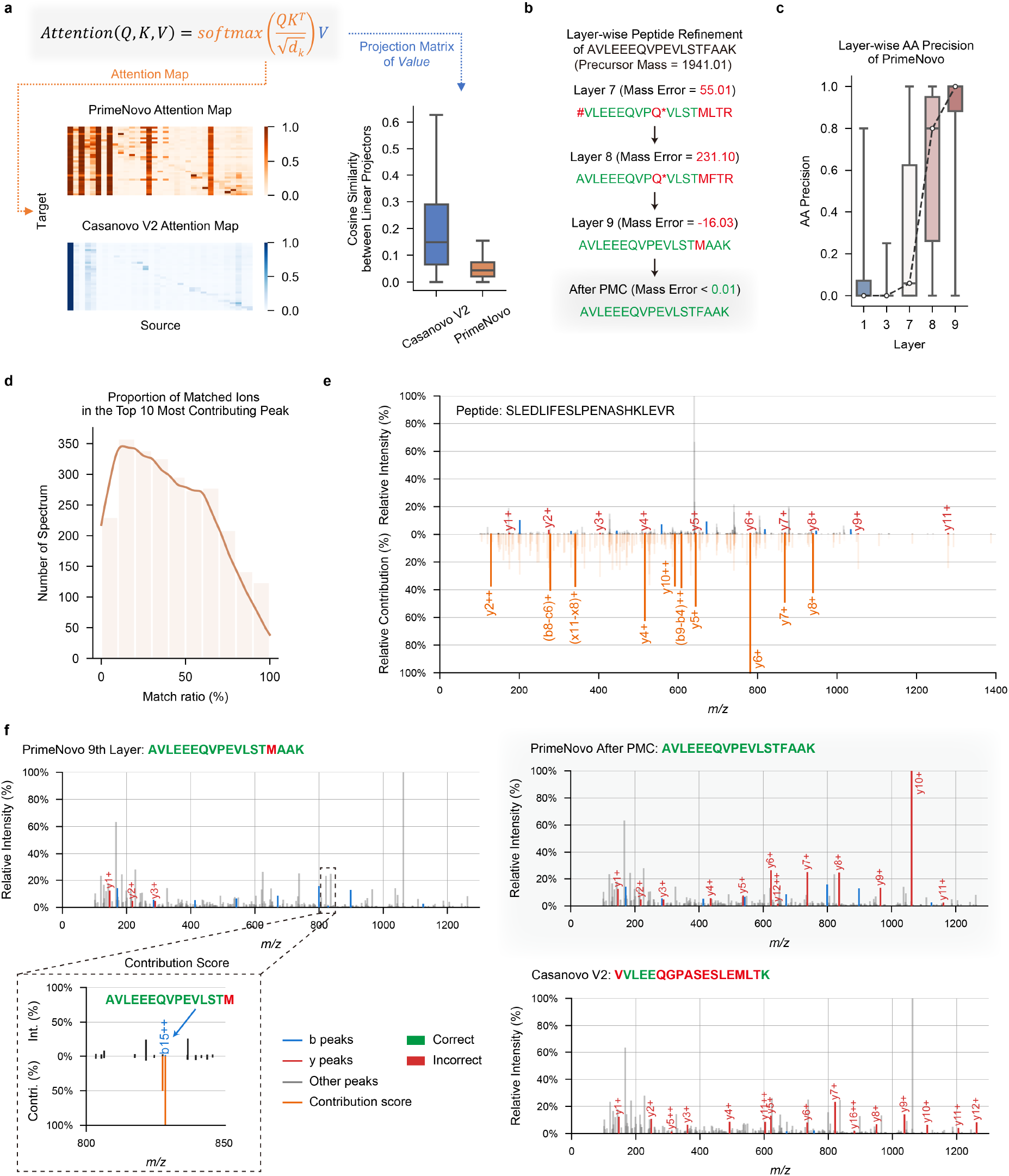
Error analysis and model explainability offer valuable insights into the performance of PrimeNovo. Attention Map and Feature Vector Similarity: This section showcases the visualization of attention maps between the Transformer encoders of Casanovo V2 and PrimeNovo. It also includes a detailed similarity analysis of each column in the feature vector from the value matrix projection. **b**. Layerwise Prediction Refinement: A case study demonstrates how PrimeNovo’s non-autoregressive model progressively refines predictions layer by layer, highlighting the model’s capacity for self-correcting its predictions as a whole. Note that * represents the Glutamine deamidation modification on amino acid Q. **c**. The chart displays the average prediction accuracy at the amino acid level across each layer in PrimeNovo. **d**. This diagram illustrates the proportion of peaks corresponding to b-y ions, as determined from predictions, based on all peaks within the PT test set ranked within the top 10 by their contribution scores. **e**. Alignment between the model’s contribution scores and the theoretical b-y ion peaks derived from predictions is presented. The diagram’s lower half shows the magnitude of all contribution scores, emphasizing those matching the b-y ions. The upper half provides a comparison with the original spectrum. **f**. A case study on how the theoretical ions, calculated from the predicted peptide, align with the input spectrum. The matched theoretical b-y ions are distinctly marked in red and blue for predictions made by PrimeNovo and Casanovo, respectively. This comparison seeks to identify potential sources of error in incorrect predictions. The diagram’s bottom left section highlights a high contribution score assigned to an incorrect peak, corresponding to a b-ion peak linked to an erroneous amino acid prediction in PrimeNovo’s final layer.

Furthermore, we conducted a numerical comparison of the Value matrices learned by the encoder networks of both models [30]. Each column in the Value matrix projection represents a hidden feature. To assess the diversity of features present in the Value matrix, we calculated the average cosine similarity between every pair of columns. As illustrated in the bar plot in Fig. 4a, it is evident that PrimeNovo’s feature vectors exhibit lower similarity to each other, as indicated by the lower average cosine similarity values in the plot. This suggests that our model’s Value matrix encompasses a broader spectrum of information and a more diverse set of features [31, 32]. This finding could provide an additional explanation for our model’s superior performance. For a more comprehensive assessment of the orthogonality of the Value matrix projection, which is evaluated by measuring the norm of the Gram matrix [30–32] (see the Supplementary Fig. 14).

Since our non-autoregressive model predicts the entire sequence at once, we can examine how each of the nine model layers progressively improves the overall sequence prediction. We decode the whole sequence from each layer of our model and observe how the amino acids evolve over time. As illustrated in Fig. 4c, amino acid-level accuracy experiences a significant surge from layer seven to nine, with a consistent increasing trend across each layer. This signifies a continual improvement in prediction accuracy at each layer. By examining the case study presented in Fig. 4b, we discern that this increase in accuracy is achieved through a layer-wise self-correction mechanism. In this process, each layer gradually adjusts the erroneously predicted amino acids throughout the entire sequence, making them more reasonable and closer to the true answer. The non-autoregressive model’s capability of enabling each amino acid to reference the surrounding amino acids for information facilitates accurate and effective correction across its layers. PMC, acting as the final safeguard against errors, rectifies model prediction errors by selecting the most probable sequence that adheres to the mass constraint. This process yields a slightly modified sequence compared to the output from the last layer, ultimately leading to the correct answer.

We also employed the feature contribution technique saliency maps [33] to analyze the impact of each peak on the prediction results. This technique generates contribution scores that provide a quick view of the impact of each peak on the prediction. A higher contribution score for a peak indicates a larger impact on the results. On the test set of PT, the contribution scores for all peaks in each spectrum. Subsequently are calculated, we sorted all peaks in descending order based on their contribution scores and selected the top 10 peaks. Using the known peptide sequences associated with these spectra, all possible fragmentions considering only 1+ and 2+ ions are generated using the in-house script (see Supplementary Section 5 for more details). We then compared the *m/z* values of the top 10 peaks with the *m/z* values of all possible fragment ions, considering a match if the difference was within 0.05 Da. Finally, the percentage of the top 10 peaks that could be matched is calculated. As shown in Fig. 4d, approximately 40% of the spectra had a matched percentage of above 50%. Importantly, our model not only focused on the major peaks but also considered intermediate fragment ions. For example (Fig. 4e), in the spectrum corresponding to the peptide sequence “SLEDLIFESLPENASHKLEVR”, among the top 10 peaks with the highest contribution scores, seven were *b* ions, while the remaining three corresponded to intermediate fragment ions “FE ((*b*8 − *c*6)+)”, “LIFES ((*b*9 − *b*4)+)”, and “PEN ((*x*11 − *x*8)+)”, respectively. These results demonstrate that our model learned a few informative peaks from the spectra, which are useful for peptide inference.

To analyze which peak in the spectrum led to the erroneous generation of the model, we visualized the spectrum by highlighting *b*-*y* ion peaks corresponding to the model’s predictions. As shown in Fig. 4f, Casanovo V2’s predicted sequence predominantly aligns its *y*-ions with input spectrum peaks, with very few calculated *b*-ions aligning with input peaks. This behavior is a consequence of the autoregressive model’s prediction direction from right to left, making it more natural to choose *y*-ion peaks for forming predictions. However, given the presence of noise in the spectrum, this prediction approach can lead to errors when *y*-ions are inaccurately selected, as demonstrated in Fig. 4f. In contrast, PrimeNovo’s predictions exhibit an alignment with both *b*-ions and *y*-ions in the input spectrum. This is due to our model’s prediction process, which leverages information from both directions, allowing it to effectively utilize the peak information from both ends of the sequence. Furthermore, we conducted a detailed analysis to identify the specific peak responsible for prediction errors in the last layer. This is achieved by calculating a gradient-based contribution score for each input peak, serving as a robust indicator of which input has a greater impact on the output, determined by the magnitude of the gradient. As observed in the left corner of Fig. 4f, the highest contribution scores across the entire spectrum coincide precisely with the peak corresponding to PrimeNovo’s incorrectly predicted *b*-ion, and this critical information is captured and corrected by our PMC unit.

### PrimeNovo demonstrates exceptional performance in taxon-resolved peptide annotation, enhancing metaproteomic research

We conducted an evaluation to gauge PrimeNovo’s proficiency in enhancing the identification of taxon-unique peptides, particularly in the context of metaproteomic research. The field of metaproteomics poses significant challenges when it comes to taxonomic annotation, primarily due to the vast diversity within microbiomes and the presence of closely related species that share high protein sequence similarity. Consequently, increasing the number of unique peptides represents a crucial approach for achieving precision in taxonomic annotations. In our assessment, we turned to a metaproteomic dataset [34] obtained from gnotobiotic mice, hosting a consortium of 17 pre-defined bacterial strains (as summarized in Supplementary Table 2). Within this dataset, we applied PrimeNovo and Casanovo V2 ^2^ to sequence unidentified MS/MS spectra through database search, all without the need for fine-tuning [34].

As illustrated in Fig. 5a, PrimeNovo exhibits superior performance compared to Casanovo V2, identifying a significantly higher number of PSMs (8,446 vs 4,072) and peptides (3,157 vs 1,412) following the rigorous quality control process T\U\D\DS, resulting in a relative increase of 107% and 124%, respectively. Furthermore, PrimeNovo excels in enhancing taxonomic resolution, outperforming Casanovo V2 in the detection of taxon-specific peptides. Notable increases are observed in bacterial-specific (1,047 vs 520), phylum-specific (828 vs 399), genus-specific (511 vs 241), and species-specific (215 vs 92) peptides (Fig. 5b-d). Particularly noteworthy is the high identification accuracy achieved by PrimeNovo, where all identified peptides are correctly matched to known species, while Casanovo V2 exhibits one incorrect matching at the genus level (Fig. 5c).

**Fig. 5.**
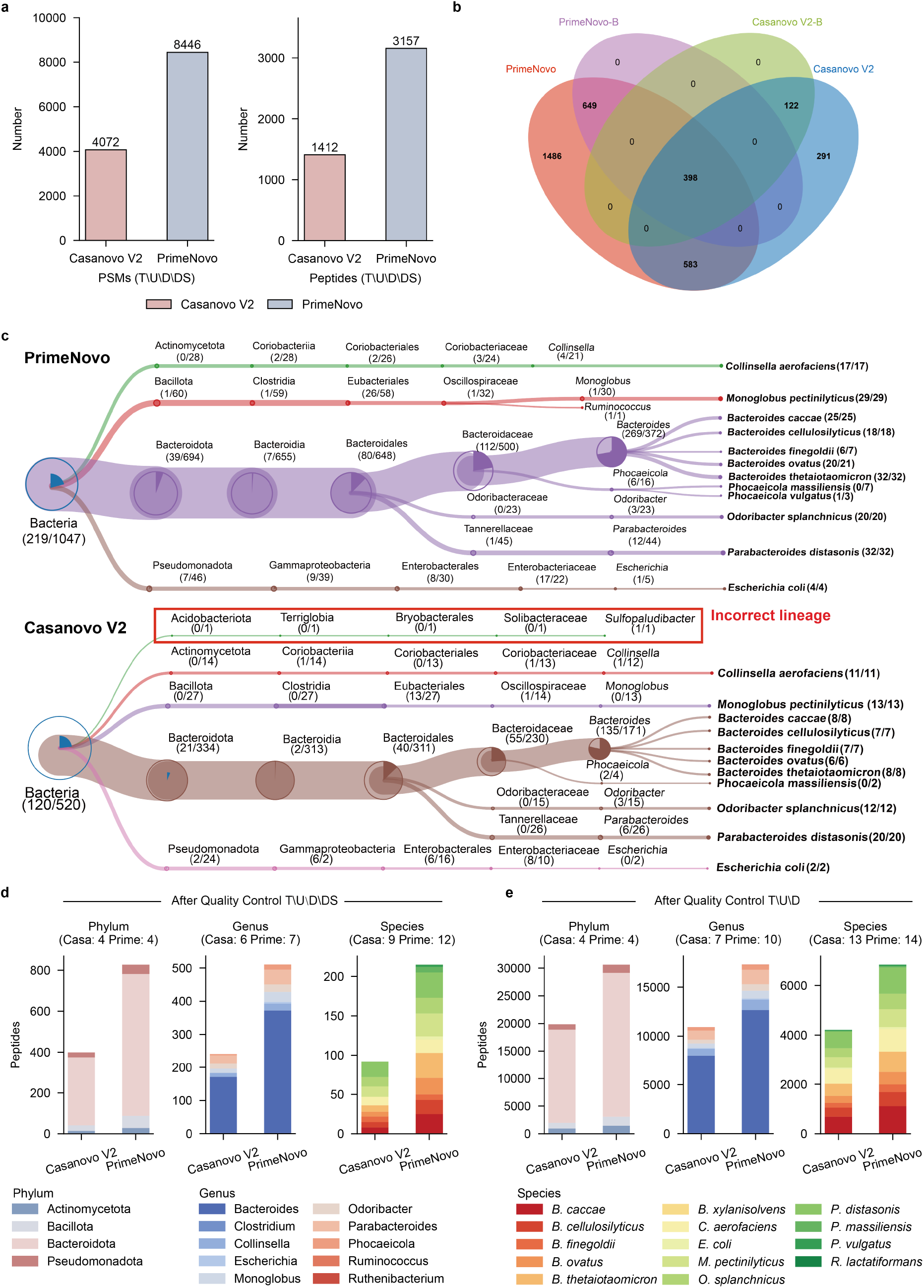
The advantages of PrimeNovo in metaproteomic analysis. **a**. Identification of PSMs and peptides through the quality control process T*\*U*\*D*\*DS. **b**. The Venn diagram illustrates the overlap between peptides identified by Pri-meNovo and Casanovo V2, as well as the bacterial-specific peptides (PrimeNovo-B and Casanovo V2-B). **c**. The treeview representation of species-level identification. **d**. The number of peptides identified at the phylum, genus, and species levels, with the note that taxa identified by fewer than three unique peptides are excluded. **e**. The number of peptides at the phylum, genus, and species levels after the quality control process T*\*U*\*D.

We further conducted an analysis of high-confidence identification results under the quality control process T\U\D. PrimeNovo demonstrated a significant increase in both PSM and peptide identifications, with a 66% increase (513,590 vs 308,499) in PSMs and a 46% increase (58,392 vs 39,866) in peptides. This result is further emphasized by the higher identifications of taxon-unique peptides achieved by PrimeNovo, surpassing Casanovo V2 in several categories, including bacterial-specific (36,704 vs 24,349), phylum-specific (30,652 vs 19,866), genus-specific (17,332 vs 10,906), and species-specific (6,848 vs 4,209) peptides (Fig. 5e). Subsequently, we assessed the models’ performance in taxonomic annotation at the protein level, which is crucial for enhancing the taxonomic resolution and contributing to subsequent research in the taxon-function network. As depicted in Fig. 5f, proteins identified by PrimeNovo and Casanovo were correctly assigned to 10 genera, 14 species, and 20 COG (Clusters of Orthologous Groups of proteins) categories. On the genus level, PrimeNovo identified a total of 6,883 proteins assigned to the 10 genera, with 6,709 of them annotated to specific COG functions. In contrast, Casanovo V2 identified only 5,028 proteins, with 4,896 of them annotated. Thus, PrimeNovo achieved a 36.89% and 37.03% increase over Casanovo V2 in taxon and functional annotations.

Furthermore, a detailed examination at the genus level revealed that PrimeNovo increased the number of proteins assigned to each genus compared to Casanovo V2: Bacteroides (4,926 vs 3,623), Clostridium (3 vs 2), Collinsella (486 vs 383), Escherichia (91 vs 62), Monoglobus (294 vs 197), Odoribacter (297 vs 204), Parabacteroides (576 vs 425), Phocaeicola (204 vs 130), Ruminococcus (3 vs 1), Ruthenibacterium (3 vs 1). Similarly, PrimeNovo exhibited significant potential for taxonomic annotation at the species level. Compared to Casanovo V2, PrimeNovo identified an additional 45.32% (3,136 vs 2,158) proteins assigned to the 14 species, with 45.03% (3,034 vs 2,092) of these proteins annotated to specific COG functions. These results demonstrate that PrimeNovo significantly enhances taxonomic resolution at both the peptide and protein levels, highlighting its substantial potential in metaproteomic research.

### PrimeNovo enables accurate prediction of a wide range of different post-translation modifications

PTMs play a crucial role in expanding the functional diversity of the proteome [35], going well beyond the inherent capabilities of the genetic code. The primary challenge lies in the underrepresentation of modified peptides within the dataset, especially those that have not been enriched for certain modifications. The detection of such peptides is often overshadowed by the more prevalent unmodified peptides. Moreover, the distinct physical properties of modified residues—namely their mass and ionization efficiency—further complicate the detection [36–40]. The capabilities of current database search engines are limited, permitting the consideration of only a select few modifications. This scarcity leads to a low presence of modified peptides in the training data, thereby making it difficult for models to accurately identify diverse PTMs from spectral data.

To address these challenges, PrimeNovo has been advanced in predicting peptide sequences with multiple PTMs, establishing itself as a foundational model divergent from conventional methods that start anew for each PTM type. By fine-tuning on enriched PTM data, PrimeNovo gains extensive exposure to multiple PTM types while retaining its ability to recognize standard peptides. Architectural adjustments, as illustrated in Fig. 6a, including the addition of a classification head above the encoder to identify specific PTMs and a newly initialized linear layer above the decoder, enhance PrimeNovo’s ability to decode peptides with PTMs, broadening the model’s token repertoire. The final loss is formulated in a multi-task setting, combining the peptide decoding loss with a binary classification task for PTM identification loss.

**Fig. 6.**
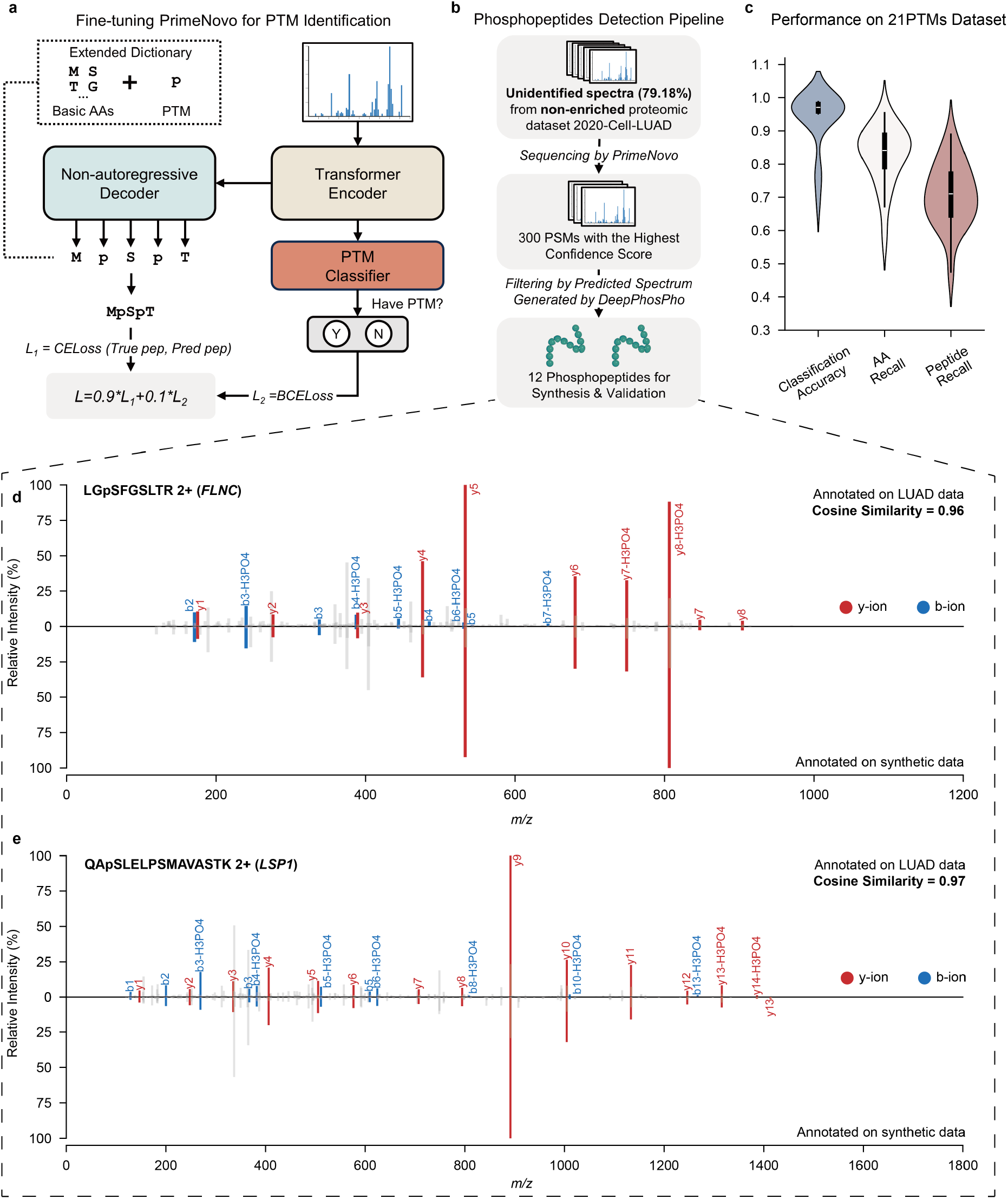
De novo sequencing of peptides with PTMs. **a**. A fine-tuning pipeline for PrimeNovo’s PTM prediction. The methodology for selecting high-quality phosphopeptides predicted by PrimeNovo. **c**. Performance metrics on the 21PTMs dataset, including classification accuracy, amino acid-level recall, and peptide-level recall. **d-e**. A comparative analysis of the actual input spectrum and the spectrum of the synthesized peptide predicted by PrimeNovo. The diagrams’ upper sections display the original input spectrum, whereas the lower sections illustrate the spectrum generated from the predicted peptide sequence. Overlapping peaks are highlighted in red and blue for b-y ions. The Cosine Similarity is calculated based on spectrum encoding using the GLEAMS package.

Our training methodology employed a dataset encompassing 21 distinct PTMs, referred to as the 21PTMs dataset, as detailed in [41]. We fine-tuned PrimeNovo for each PTM to ascertain its proficiency in peptide generation and PTM classification, in accordance with previously described methods. To ensure dataset balance, we included an approximately equal number of peptides with and without PTMs, culminating in a total of 703,606 PSMs for the dataset. The comprehensive fine-tuning endeavor across the 21 PTMs allows PrimeNovo to discern a broad spectrum of PTMs, a capability evidenced by the exemplary performance metrics for each PTM category depicted in Fig. 6c. Specifically, the classification accuracies for all PTMs exceeded 95%, except asymmetric and symmetric Dimethylation at Arginine (R), and Monomethylation at Arginine (R), which have classification accuracies of 77%, 77%, and 69%, respectively. Excluding Monomethylation at Arginine (R), which recorded a peptide recall rate of 48%, the de novo sequencing recall for peptides with the other 20 PTMs exceeded 61%. Such peptide recall levels are on par with performance in other datasets without special PTMs, such as an average peptide recall of 64% across nine-species datasets. Detailed insights into the classification accuracy and peptide recall for each PTM are provided in the supplementary Fig. 16.

To assess PrimeNovo’s inference performance on PTMs within a more applied context, we selected a phosphorylation dataset from Xu et al. [42] (denote as the 2020-Cell-LUAD dataset), which concentrates on Human Lung Adenocarcinoma with 103 LUAD tumors and their corresponding non-cancerous adjacent tissues. It offers both phosphorylation-enriched and non-enriched data. We randomly selected a portion (3389 PSMs) of the enriched data for testing and the rest for training, checking of no overlapping peptide sequence between the training and testing sets. We fine-tuned PrimeNovo on such training data and the test results demonstrate that PrimeNovo distinguishes between phosphorylated and non-phosphorylated spectra with a classification accuracy of 98%, and achieves a peptide recall rate of 66% on both cancer tissue data and non-cancerous adjacent tissues test data, as detailed in Supplementary Table 9. To assess PrimeNovo’s capability to identify modified peptides within non-enriched proteomic datasets, we deployed it for the analysis of unidentified MS/MS spectra from the non-enriched 2020-Cell-LUAD dataset, notably without conducting dataset-specific fine-tuning. Given the absence of peptide identifications from existing databases in this dataset, we relied on the model’s confidence scores to select 300 high-quality predicted peptides. We then undertook a comparative analysis between the theoretical spectrum, as generated by DeepPhosPho [43], and the original input spectrum corresponding to these peptides, as illustrated in Fig.6b. Through this process, we pinpointed 12 peptides as candidates for synthesis validation and further functional investigation. The details of the selection methodology are elaborated upon in Supplementary Section 7.

All 12 phosphopeptides predicted by PrimeNovo from non-enriched data were validated using their synthetic counterparts, as depicted in Fig. 6 and Supplementary Fig. 17 and Supplementary Fig. 18. In Fig. 6. d,e, they showcase the alignment between theoretical and experimental spectra for two representatives of 12 synthesized phosphorylated peptides. The comparison reveals a strong correspondence between the predicted b-ions and y-ions peaks and the experimental spectrum’s signal peaks, evidenced by a Pearson correlation exceeding 0.90 for nine paired spectra, and 0.70, 0.72, and 0.86 for the remaining three pairs. This correlation underscores the model’s high predictive precision. Further investigation into the proteins associated with these phosphopeptides highlighted their relevance to lung adenocarcinoma (LUAD). For example, the peptide LGpSGFSLTR (2+) (Fig. 6d) from Filamin-C (FLNC) aligns with findings that the ITPKA and Filamin C interaction fosters a dense F-actin network, enhancing LUAD cell migration [44]. Another identified peptide, HGpSDPAFAPGPR (2+) from FAM83H (Fig. 6e), is noted for being upregulated in LUAD, indicating a potential prognostic marker of LUAD [45, 46]. Additionally, peptides WLDEpSDAEMELR, GPAGEAGApSPPVR, and AQpTPPGPSLSGSK reveal proteins (HACD3, SNTB2, and SRRM2) not previously associated with LUAD, but there are studies suggesting potential relevance between these three proteins and other cancer types. This offers new directions for potential biological research on the disease by examining the above-relevant proteins. For detailed results concerning the remaining peptides and the comprehensive experimental methodologies used for their synthesis and analysis, please see Supplementary Section 6.

These results demonstrate that PrimeNovo has a high sensitivity in detecting PTMs from proteomic datasets, especially those non-enriched ones, which provides a novel solution for low-abundance PTM discovery.

## Discussion

Peptide sequencing is essential for deciphering protein structures and functions. In this work, we introduce PrimeNovo, a Transformer-based deep learning model designed to perform rapid and accurate de novo peptide sequencing tasks. Informed by deep biological insights, PrimeNovo utilizes a non-autoregressive architecture for efficient bidirectional peptide sequencing and a unique PMC decoding unit for precise, mass-specific controllable generation. PrimeNovo has consistently demonstrated exceptional accuracy at both peptide and amino acid levels, as evidenced by its state-of-the-art performance in multiple spectrum test datasets. The significant speedup we have achieved makes it highly suitable for large-scale peptide sequencing tasks. Moreover, Our model exhibits robust adaptability to a wide range of data sources and distributions, whether under zero-shot or fine-tuning settings. This versatility allows our model to be seamlessly deployed in various applications while maintaining excellent performance. In practical downstream applications, PrimeNovo excels in peptide annotation for metaproteomic research, making it easier to identify specific microorganisms within the given samples and elucidate their functional roles. PrimeNovo’s ability to handle newly added amino acid tokens enables accurate PTM detection. This feature has been instrumental in identifying numerous potential new peptides that may not have been detected using traditional database search algorithms.

## Methods

### Datasets

The primary dataset used for training our model is the MassIVE Knowledge Base spectral library version 1 (MassIVE-KB) [26], which we obtained from the MassIVE repository. This extensive dataset comprises over 2.1 million precursors originating from 19,610 proteins. These precursors were distilled from a vast pool of human data, amounting to more than 31 terabytes, gathered from 227 public proteomics datasets within the MassIVE repository.

To ensure a fair and comprehensive evaluation of our model’s performance, we incorporated various datasets into our study. We acquired the widely used nine-species benchmark dataset [12] for benchmarking purposes. Subsequently, we conducted thorough assessments on several datasets, including the synthesized Proteometools (PT) dataset, the real-world human hepatocellular carcinoma proteomics dataset (HCC), the multi-enzyme antibody dataset IgG1-human-HC, and the revised nine-species benchmark datasets. In addition, to compare our model with the recently introduced CNN-based approach PepNet [19], we obtained their test dataset, PXD019483 [47]. Furthermore, to assess our model’s performance in comparison with the graph-based approach GraphNovo [23], we acquired their test dataset, which encompasses samples from *A. thaliana, C. elegans*, and *E. coli*, referred to as the three-species dataset. A summary of the training and testing datasets used in this study can be found in Supplementary Table 1.

To evaluate PrimeNovo’s ability to enhance the identification of taxon-unique peptides in metaproteomic research, we obtained a publicly available human-gut-derived bacterial metaproteomics dataset [34]. Additionally, to assess our model’s performance in analyzing peptides with PTMs, we obtained two datasets. One is the publicly available benchmark dataset 21PTMs [41], notable for containing the largest variety of PTMs to date. The other is a publicly available phosphoproteomics dataset [42], focused on the study of lung adenocarcinoma (LUAD). Further details about these datasets are provided in the “Datasets” section of the Supplementary Information.

### Overview & Notation

In this section, we present a comprehensive overview of the problem setting and the details of our proposed approach for peptide de novo sequencing. Our approach comprises two key components: a non-auto regressive Transformer (NAT) model [48], which is initially trained using the connectionist temporal classification (CTC) loss [49], and a dynamic programming (DP) solver that carefully selects the amino acids based on prediction probability while satisfying the precise mass constraint. Additionally, we introduce an optimization strategy for the DP solver, leveraging the power of Compute Unified Device Architecture (CUDA) programming to significantly enhance the inference speed, thereby improving overall performance.

In the de novo sequencing task, we are provided with a spectrum instance denoted as S = {*I, c, m*}, which is generated by a mass spectrometer when analyzing biological samples. Here, *I* = {(*m/z*_1_, *i*_1_), (*m/z*_2_, *i*_2_), · · ·, (*m/z*_*k*_, *i*_*k*_)} represents a set of Mass-to-Charge ratio and corresponding intensity pairs. These pairs are retained after being filtered by the mass spectrometer threshold. Additionally, *c* denotes the measured charge of the peptide (precursor), and *m* represents the measured total mass of this peptide. Our primary objective in this context is to derive the correct amino acid sequence denoted as *A* = {*a*_1_, *a*_2_, · · ·, *a*_*n*_} from the information contained within 𝒮.

### Non-autoregressive Transformer backbone

We adopt the Transformer encoder-decoder network as our foundational model, following the work of Casanovo [16]. In the encoder network, we handle the Mass-to-Charge ratio *m/z* and the intensity information *i* from the set *I* separately before merging them. To represent each *m/z* value, we employ a sinusoidal embedding function, which effectively captures the relative magnitude—an essential factor in determining the peptide fragments:

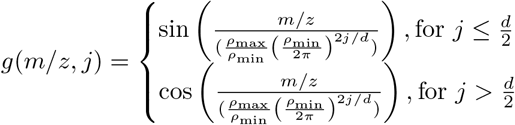

Here, *j* signifies the position in the *d*-dimensional hidden embedding. The parameters *ρ*_max_ and *ρ*_min_ define the wavelength range for this embedding. In contrast, we handle intensity values through a linear projection layer.

In the non-autoregressive model, the only architectural distinction between the encoder and decoder lies in the cross-attention mechanism. Therefore, we employ identical notations for both components. In a formal sense, each layer computes a representation *R*, based on the preceding feature embeddings. For the *k*-th layer, the representation is

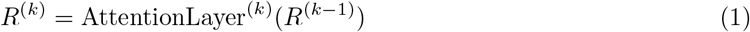

Here, *R*^(0)^ signifies the spectrum embedding for the encoder, while for the decoder, it represents the summation of positional and precursor embeddings. To maintain consistency, we keep the generation length fixed as **t** for the decoder. Consequently, the output of the final decoder layer undergoes a softmax operation, which calculates the probability distribution over tokens for each position.

### Peptide reduction strategy for our non-autoregressive modeling

The conventional approach to sequence generation involves training a system to predict the probability of the next token, denoted as *P* (*a*_(*i*+1)_|*a*_1:*i*_). This method is known as autoregressive generation, where the likelihood of the (*i* + 1)-th amino acid is conditioned on the preceding *i* amino acids. However, this autoregressive approach imposes a unidirectional flow in the generation process, preventing the system from revising previous outputs. Such a unidirectional flow is at odds with the inherent nature of proteins, where the presence of each amino acid depends on information from both preceding and succeeding amino acids. To overcome this limitation, we propose a non-autoregressive approach to sequence modeling. In this paradigm, all amino acids can be generated simultaneously. This means that each amino acid’s generation is not solely reliant on the preceding ones; instead, it can access bidirectional information from the surrounding amino acids. This better aligns with the natural behavior of proteins and significantly enhances the accuracy of our sequence generation system. Formally, in a non-autoregressive system with a predefined maximum generation length *t*, we model the probability of generating an amino acid, *P* (*a*), at each position *t* independently.

Nonetheless, the straightforward approach of modeling probabilities at each position can result in weak global connections, making the entire sequence appear nonsensical, even if regional generation is accurate. For example, during training, a phrase like “au revoir” might be translated to both “goodbye” and “see you,” leading an inference process in a non-autoregressive translation system trained with crossentropy loss to produce the ambiguous phrase “see bye” due to the absence of global coherence. To address this challenge, we opt for the CTC loss [49]. Unlike cross-entropy loss, which calculates losses at the amino acid level, CTC loss operates at a sequence level. This approach significantly enhances global connections and results in the generation of accurate peptides as a whole, mitigating issues related to local ambiguities and ensuring more coherent sequences.

To handle situations where the total number of generated tokens, represented by the maximum token length *t*, exceeds the target sequence length, we introduce a reduction function denoted as Γ(·) in non-autoregressive generation. This function merges consecutive identical amino acids, as illustrated below:

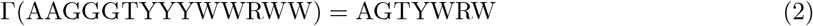

However, the straightforward reduction approach may not be suitable for peptide sequences with consecutive identical amino acids. To address this, inspired by Graves et al. (2006) [49], we introduce a special blank token denoted as *ϵ* during the generation process. Consecutive identical amino acids on both sides of *ϵ* are not merged, and *ϵ* is eventually removed, leading to the following transformation:

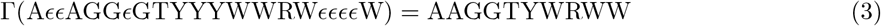

For a visual representation of this process, please refer to the Supplementary Fig. 1.

### Definition of CTC loss

Following the CTC reduction rule described above, it’s possible to obtain multiple decoding paths denoted as **y**, which can all be reduced to the target sequence *A*. For instance, both “CCGT” and “CG*ϵ*T,” among many others, can be transformed into the target sequence “CGT”. Consequently, the probability of generating the target sequence *A* is the sum of the probabilities associated with all paths **y** that can be reduced to *A*:

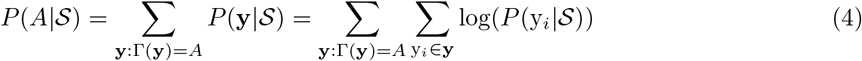

Here, **y** = (y_1_, y_2_, · · ·, y_**t**_) represents a single decoding path in the non-autoregressive model output, satisfying the condition Γ(**y**) = *A*. The overall probability of generating the target sequence *A*, denoted as *P* (*A*|𝒮), is then computed as the sum of the probabilities of generating each **y**, with y_*i*_ at each position. Since the probability is modeled independently, the probability of each **y** can be calculated as the multiplication of the probabilities of generating all y_*i*_ ∈ **y**. This multiplication can be expressed as the sum of the logarithm of the probabilities of each y_*i*_.

During the training process, our objective is to maximize the total probability of generating the target sequence *A* for each input spectrum *S*. Since we are utilizing gradient descent to optimize our model, this goal is equivalent to minimizing the negative total probability. Therefore, our loss function is simply defined as:

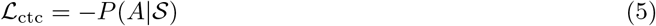

One could theoretically enumerate all possible paths **y** for each target sequence *A* in order to calculate the total probability (loss) for training our network. However, this approach becomes impractical as the number of paths grows exponentially with respect to the maximum generation length. This would result in an unmanageable amount of computation time. Instead, we adopt a dynamic programming method, as detailed in the Supplementary Information, to optimize the calculation of this loss efficiently. This approach allows us to train our model effectively w ithout t he c omputational b urden o f exhaustively enumerating all possible paths.

### Knapsack-like dynamic programming decoding algorithm for precise mass control

The generated de novo peptide sequence should be strictly grounded by molecular mass measured by the mass spectrometer. Specifically, the molecular mass of the ground truth peptide, m_*tr*_ falls in the range of [*m*−*σ, m*+*σ*], where *m* is precursor mass given by mass spectrometer, and *σ* is measurement error, usually at 10^*−*3^ level, of used mass spectrometer. However, neural network models are of low explainability and controllability, making it difficult to control the generated results to cater to certain desires. To allow the accurate generation, we reformulate the non-auto regressive generation as a knapsack-like optimization problem [50], where we are picking items (amino acids) to fill the bag with certain weight constraint, while the value (predicted log probability) is maximized. Such optimization problem can be formulated as :

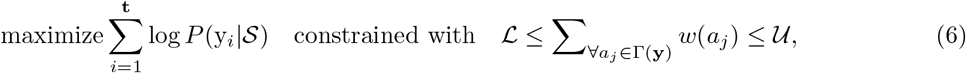

where ℒ and 𝒰 are the desired lower bound and upper bound for decoded peptide mass. We denote ℒ = m − *tol* and 𝒰 = m + *tol* where *tol* is decoding tolerance within which we think the true mass m_*tr*_ falls in, after taking into measurement error.

Inspired by a similar idea by [51], we propose a dynamic programming method to solve such an optimization task. We denote *e* as the decoding precision to construct a two-dimensional DP table. For each time step, we would have ⌈𝒰 */e*⌉ cells with being the ceiling function. The *l*-th cell can only store the peptide with mass precisely within [*e* * (*l* − 1), *e* * *l*]. Specifically, the *l*-th cell at *τ* -th time step **d**^*τ*,*l*^ stores the most probable, calculated by the sum of log probability by non-autoregressive model, *τ* tokens sequence **y**_1:*τ*_ satisfying the mass constraint of 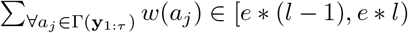.

We first initialize our DP table by filling the first time step, *τ* = 1, as follows:

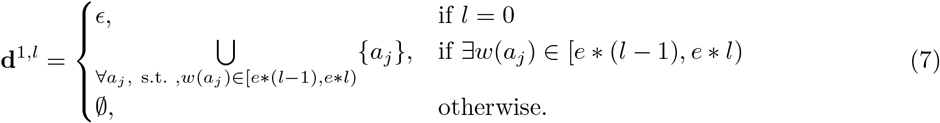

In the first case, **d**^1,1^ stores the one-token sequence with the total mass in the range of [0, *e*], where *e* is usually a very small number (*e <* 1) for higher decoding accuracy, therefore no amino acid other than *ϵ* can fall under this mass limit. On the other hand, when *l* ≠ 1, there might be multiple amino acids whose mass falls within [*e* * (*l* − 1), *e* * *l*). We store all of them in *l*-th cell to avoid overlooking of any possible starting amino acid.

We then divide the recursion steps into three cases,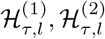 and 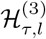, each storing its corresponding set of sequences following the rules below:

(1) When *y*_*τ*_ = *ϵ*, we know Γ(**y**_1:*τ−*1_) = Γ(**y**_1:*τ*_) due to CTC reduction, therefore the mass stay the same. This gives the set of candidate sequences :

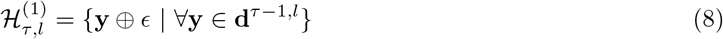

where ⊕ is the concatenation.
(2) When the newly decoded non-*ϵ* token is the repetition of the last token, the reduced sequence still remains the same with the mass unchanged, due to the CTC rule. We get the second set of potential sequences:

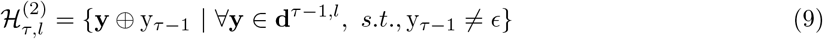
(3) When the newly decoded non-*ϵ* token is different from the last token in the already generated sequence, the mass will be increased. We select the potential sequence by examining the total mass that falls in the mass constraint:

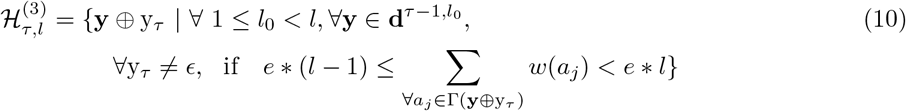 The we update the cell **d**^*τ*,*l*^ using all candidates from above three sets:

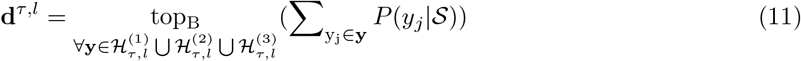

where top_B_ is taking the top B most probable sequences according to generated probability. We then select the most probable sequence at **d**^**t**,|*A*|^ cell as our final result.

### CUDA acceleration for proposed mass control decoding algorithm

The time complexity of our proposed mass control dynamic programming algorithm when executed sequentially is O(*N*_*a*_ * **t** * (*U/e*)^2^), where *N*_*a*_ represents the total number of tokens (which, in our case, corresponds to the number of amino acids plus one). This complexity arises because our recursion step for each **d**^*τ*,*l*^ involves visiting all *l*_0_ cells within the range 1 ≤ *l*_0_ *< l*, while iteratively adding each amino acid to verify the total mass.

However, it’s important to note that this calculation doesn’t take into account the potential for parallelism within the recursion process. Specifically, the update of each **d**^*τ*,*l*^ relies solely on cells from the previous time step, *τ* − 1, and can thus be parallelized along the *l* direction. Furthermore, for a given cell **d**^*τ*,*l*^ with a fixed amino acid *a*_*i*_, we only need to consider cells from 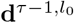, where *l*_0_ falls within the range defined by the inequality:

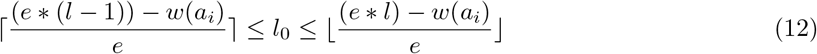

This optimization saves us from visiting all *l* cells for each amino acid, significantly improving efficiency. To implement the parallel algorithm for the PMC unit, we employ the Compute Unified Device Architecture (CUDA). CUDA is a parallel computing programming framework developed by NVIDIA, which allows programs to leverage the computational power of NVIDIA GPUs (Graphics Processing Units) for a wide range of general-purpose computing tasks. Detailed information regarding our CUDA algorithm is provided in the Supplementary Information.

### Data availability

The nine-species benchmark dataset [12] was directly downloaded as Mascot Generic Format (MGF) files from the Mass Spectrometry Interactive Virtual Environment (MassIVE) repository (identifier: MSV000081382), shared by the authors of the DeepNovo paper. The dataset was searched using PEAKS DB [5] software [version 8.0] with a false discovery rate (FDR) of 1%. The MassIVE-KB dataset [26] was obtained by downloading the Raw files and the filtered identification results from the “All Candidate library spectra” section of the MassIVE Knowledge Base spectral library v1 (https://massive.ucsd.edu/ProteoSAFe/static/massive-kb-libraries.jsp). The PT [28], 21PTMs [41] and PXD019483 [47] datasets were obtained by downloading the Raw files and MaxQuant [2] identification results from the PRIDE [52] repository by PXD004732 [28], PXD009449 [41] and PXD019483 [47], respectively. The HCC [29] and 2020-Cell-LUAD [42] datasets were obtained by downloading the Raw files and MaxQuant identification results from the iProX [53] repository (identifier: IPX0000937000 and IPX0001804000, respectively). The IgG1-Human-HC [27] dataset was obtained by downloading the combined identification results of the database algorithms MS-GF+ [54] and X!Tandem [55] with an FDR rate of 1% from the MassIVE repository (identifier: MSV000079801). The three-species dataset [23] was obtained by downloading the SEQUEST [3] search results with a 1% false positive rate for these three species datasets from the data shared by the GraphNovo authors on Zenodo (identifier: zenodo.8000316). The revised nine-species benchmark dataset [16] was obtained by downloading the Raw files and Crux [56] identification results from the MassIVE repository (identifier: MSV000090982). The Cell-metaproteome dataset [34] was obtained by downloading the Raw files and MyriMatch [57] identification results from the MassIVE repository (identifier: MSV000082287).

## Supplementary Information for

### 1 Datasets

**Supplementary Table 1:**
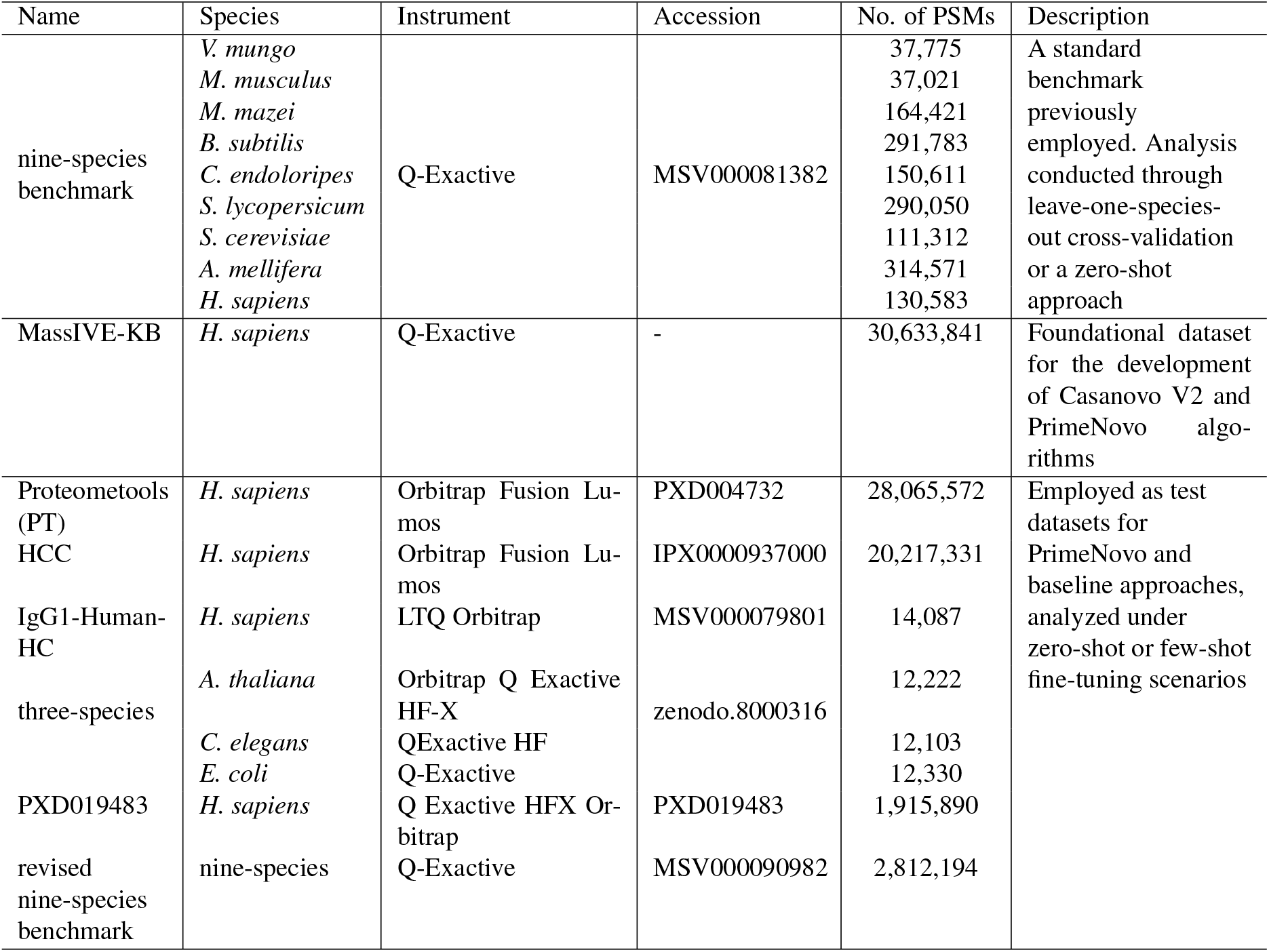
Summary of training and testing datasets in our study.

Supplementary Table 1 provides a comprehensive overview of the training and testing datasets utilized in our study. This includes the Massive-KB dataset, which was used for training PrimeNovo, and the nine-species benchmark dataset, employed for training PrimeNovo CV and for conducting comparative evaluations against baseline models. Following the training phase, PrimeNovo was subsequently evaluated on a diverse set of datasets, encompassing PT, HCC, IgG1-human-HC, three-species, PXD019483, and the revised nine-species benchmark datasets.

### 1. nine-species benchmark

dataset [1]: It was originally curated by Tran *et al*. during the development of DeepNovo [1], comprises a diverse assortment of data from nine distinct species. It was meticulously compiled by aggregating contributions from multiple research teams, with the aim of minimizing biases associated with species and laboratory sources. This dataset has since gained popularity as a preferred choice for evaluating various de novo sequencing algorithms, including Casanovo [2] and PointNovo [3], utilizing the leave-one-species-out cross-validation method established by DeepNovo. In our study, we accessed this dataset from the MassIVE repository (MSV000090982) and adopted the same cross-validation strategy to ensure a fair and consistent comparison.

### 2. MassIVE-KB [4]

This dataset played a pivotal role in training our model. It encompasses an extensive collection of over 2.1 million precursors derived from 19,610 proteins. This dataset was meticulously assembled, drawing from more than 31 TB of human data originating from 227 public proteomics datasets. Notably, it upholds rigorous false discovery rate controls, ensuring a comprehensive and robust training environment for our model.

### 3. ProteomeTools (PT) [5]

The PT dataset, chosen for PrimeNovo’s performance evaluation and fine-tuning, constitutes a carefully curated assembly of synthetic human peptides. As a prominent component of the Proteometools project, it comprises in excess of 330,000 synthetic tryptic peptides, encompassing a diverse spectrum of human gene products. The presence of well-established peptide sequences within this dataset serves as a reliable foundation for evaluating the algorithm’s precision and accuracy.

### 4. HCC [6]

From a real-world application standpoint, the HCC dataset, centered on proteomics data from early-stage human hepatocellular carcinoma (HCC) patients, was incorporated. This dataset provides a distinctive insight into the proteomics of both tumor and non-tumor tissues, facilitating an evaluation of our model’s generalizability in a clinical context.

### 5. IgG1-Human-HC [7]

The IgG1-Human-HC dataset, employed to assess the generalizability of de novo sequencing models, comprises human antibody sequences that have undergone processing with various enzymes, including trypsin, chymotrypsin, and others. This dataset holds particular importance due to its applicability in identifying novel or unfamiliar protein sequences, especially in the realm of immunotherapy antibodies where variable sequences are often unavailable, rendering traditional database search algorithms ineffective. Hence, we utilized this dataset to evaluate our model’s proficiency in unraveling the amino acid sequences of unknown antibodies.

### 6. three-species [8]

We employed the three-species dataset, made available by GraphNovo [8], which comprises samples from *Arabidopsis thaliana, Caenorhabditis elegans*, and *Escherichia coli*, for our comparative analysis. This dataset, subjected to trypsin digestion and analyzed using state-of-the-art mass spectrometry techniques, allowed us to conduct a thorough performance comparison between PrimeNovo and GraphNovo.

### 7. PXD019483 [9]

In order to facilitate a comparison with the recently published PepNet [10], we acquired the publicly available test dataset PXD019483, which was also used by PepNovo. This dataset, consisting of extensive human proteomics data, is accessible through the PRIDE repository under the identifier PXD019483. In line with PepNet’s approach, we filtered the spectra to include only those with charge states 2+ and 3+, as identified by MaxQuant with a false discovery rate (FDR) threshold of less than or equal to 1%, and with precursor mass differences of no more than 10 ppm.

### 8. Revised nine-species benchmark [2]

The dataset was acquired by the authors of Casanova V2 [2] through a process that involved downloading the raw files of the nine-species benchmark dataset and subsequently reanalyzing them using Crux version 4.1 [11]. Following this analysis, peptides shared between species were removed. The resulting dataset, known as the new nine-species benchmark, encompasses approximately 2.8 million PSMs and is derived from a total of 343 RAW files. For our study, we directly obtained the identification results that were shared by the authors.

For both the PT and HCC datasets, we partitioned each into separate training and test sets to facilitate further fine-tuning and evaluation. Given the substantial size of these datasets, only a minimal fraction, specifically 0.3%, was designated for testing purposes. Stringent precautions were taken to ensure the absence of any overlap in peptide sequences between the training and testing sets

Supplementary Table 2 presents a comprehensive overview of the datasets utilized for downstream applications in our investigation.

**Supplementary Table 2:** Summary of datasets used in downstream applications.

| Name | Species | Instrument | Accession | No. of PSMs | Description |
| --- | --- | --- | --- | --- | --- |
| Cell-metaproteome | Microbes | LTQ Orbitrap | MSV000082287 | 26,457,107 | For metaproteomics analysis |
| 21PTMs | *H. sapiens* | Orbitrap Fusion Lumos | PXD009449 | 703,606 | For PTM analysis |
| 2020-Cell-LUAD | *H. sapiens* | Orbitrap Fusion | IPX0001804000 | 26,162,410 |  |

### 1. Cell-metaproteome [12]

This dataset was utilized to assess the performance of our model on complex samples containing multiple coexisting species, such as microbial communities that often lack reference sequences. We accessed a publicly available human-gut-derived bacterial metaproteomics dataset, which was sampled from the human gastric organ and processed using an LTQ Orbitrap mass spectrometer, resulting in the generation of MS/MS spectra. These MS/MS spectra were subjected to analysis using MyriMatch v.2.2. We obtained approximately 26 million PSMs by downloading the identification results generously shared by the authors of the analysis [12]

### 2. 21PTMs [13]

The 21PTMs dataset represents a publicly available benchmark dataset notable for containing the most extensive variety of post-translational modifications (PTMs) to date. This dataset comprises approximately 5,000 peptides, which in turn represent 21 different naturally occurring human PTMs, encompassing modifications of lysine, arginine, proline, and tyrosine side chains, along with their corresponding unmodified counterparts.

### 3. 2020-Cell-LUAD [14]

To assess our model’s ability to infer phosphorylated peptides, we obtained a publicly available phosphoproteomics dataset, known as 2020-Cell-LUAD [14]. This dataset was specifically designed for the study of lung adenocarcinoma (LUAD) and was compiled from 103 LUAD tumors and their corresponding non-cancerous adjacent tissues. It offers both phospho-enriched data (the phosphoproteomic data) and non-enriched data (the proteomic data), facilitating a comprehensive analysis of phosphorylation events.

## 2 Additional model details

### 2.1 CTC loss calculation with dynamic programming

As we mentioned in “Section Methods. Definition of CTC loss” of our main manuscript, CTC loss hinges on calculating the total probability of all paths that lead to a desired target sequence. However, enumerating all paths in the model output to calculate the *P* (*A*|𝒮) is too time-consuming as there are (m + 1)^**t**^ different decoding paths, with m being the total number of amino acids adding one *ϵ* token. To enable efficient calculation, dynamic programming is used during the training using CTC. Let denote *α*(*τ, r*) as the probability of generating *A*_1:*r*_, the first *r* amino acids in the ground truth sequence *A*, using only the output positions up to *τ*, formally:

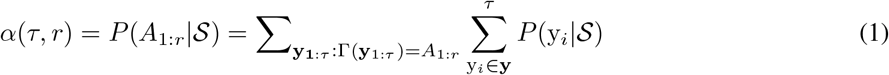

Consequently, *α*(*τ*, 0) is the probability of generating *ϵ* tokens at all first *τ* positions in the model output, as |Γ(**y**_1:*τ*_)| = 0 ⇒ **y**_1:*τ*_ = (*ϵ, ϵ*, · · ·, *ϵ*). And *α*(1, 1) is the probability of generating the exact first true amino acid at the first position. We initialize our DP as follows:

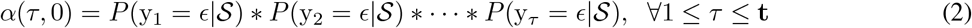

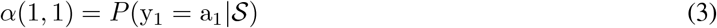

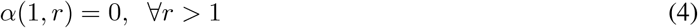

For the convenience of explanation, we further decompose the *α*(*τ, r*) into *α*(*τ, r*|y_*τ*_ = *ϵ*) + *α*(*τ, r*|y_*τ*_ ≠ *ϵ*) by the law of total probability. We then recursively calculate *α*(*τ, r*) using the following rule:

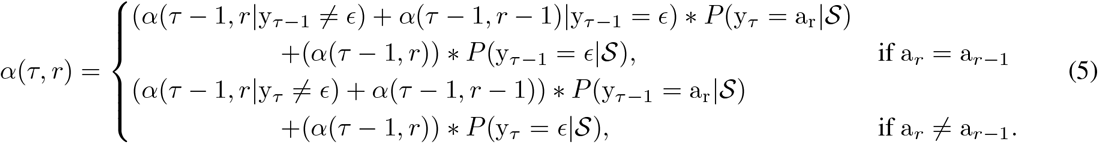

When a_*r*_ = a_*r−*1_, either by adding the *a*_*r*_ to decoded sequence up to*τ* − 1 time step **y**_*τ−*1_ which can be reduced to *A*_1:*r−*1_, with y_*τ−*1_ = *ϵ*, or repeating the last token of **y**_*τ−*1_ which can be reduced to *A*_1:*r*_ can both yield *A*_1:*r*_. Otherwise, when a_*r*_ ≠ a_*r−*1_, we wouldn’t need to consider y_*τ−*1_ = *ϵ* when adding *a*_*r*_, with everything else being the same.

Then, our loss function L_ctc_ is:

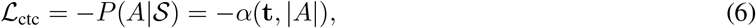

as we are trying to maximize the total probability of all paths that can be reduced to *A*. This is precisely the value of *α*(**t**, |*A*|) as we defined above. We make it negative to make it a minimization problem.

**Supplementary Fig. 1:**
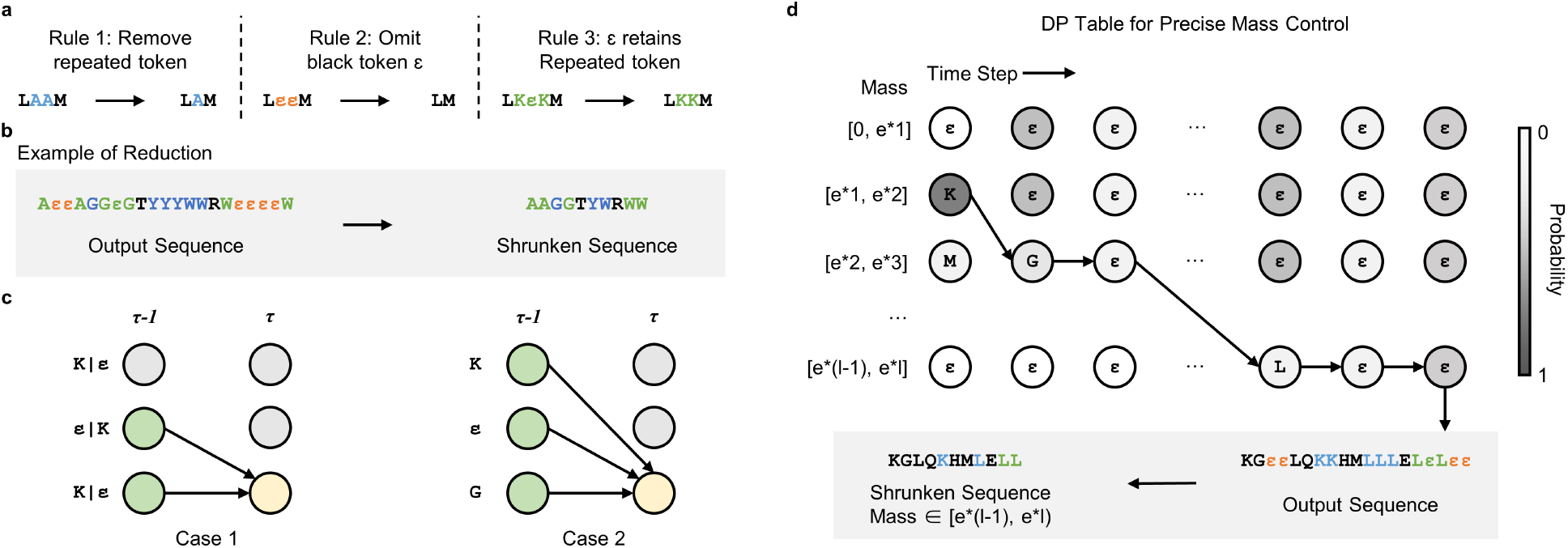
**a**. Visualization of the CTC Token Processing Reduction Rule. **b**. A case study of how the reduction rule can be applied to a sequence to obtain the final sequence. **c**. Two Cases in CTC Loss Calculation: On the left, the scenario where the upcoming token is a repeat of the previous one. On the right, the scenario where the next token differs from the preceding one **d**. Overview of the PMC Decoding Process: Sequential decoding from the first to last time step with mass constraint adherence in Each Cell, culminating in the application of the CTC Reduction Rule for the final output.

### 2.2 CUDA algorithm for PMC unit

#### Algorithm 1 CUDA PMC Algorithm.

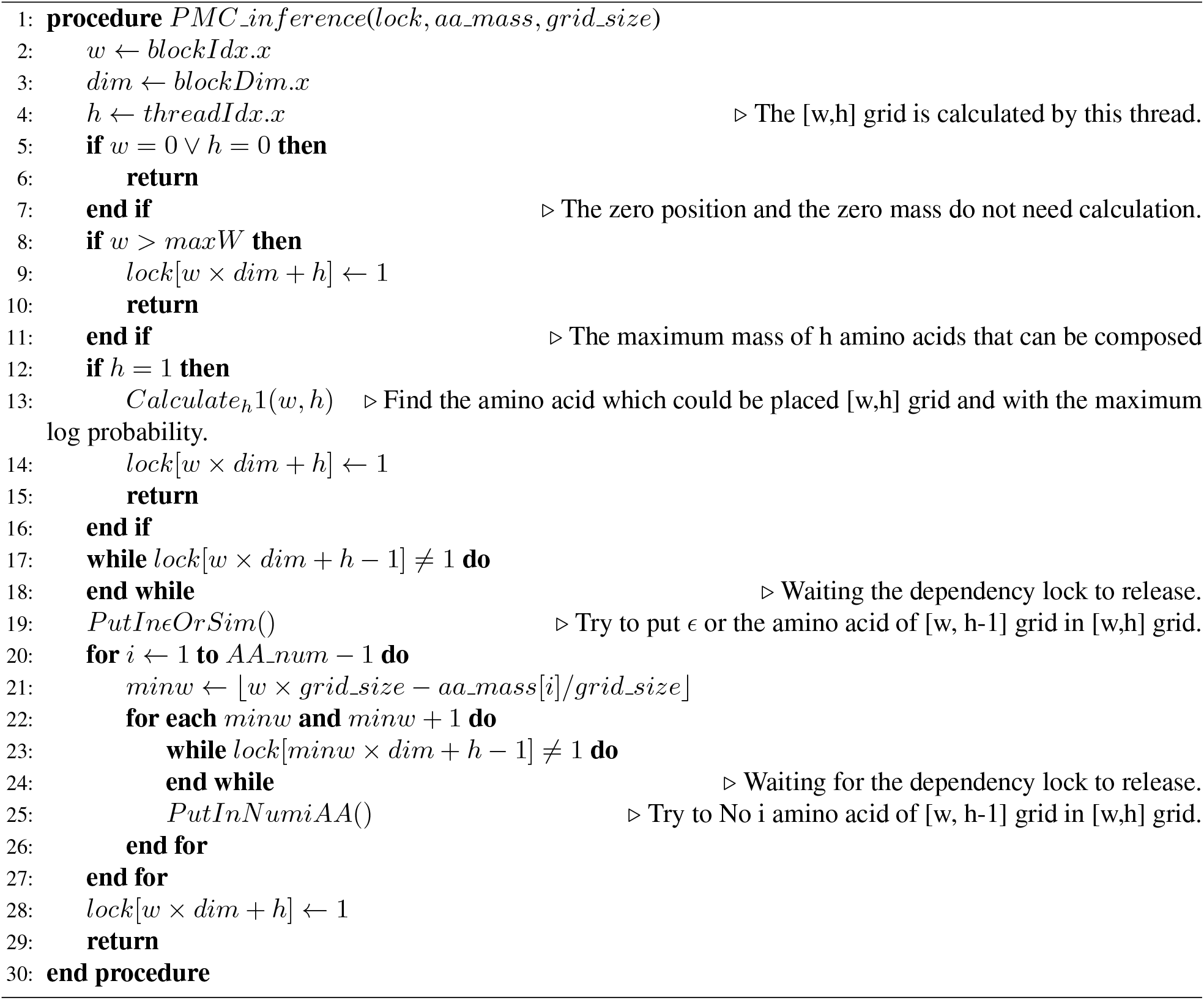

Our CUDA algorithm begins by constructing a two-dimensional dynamic table. In this table, we store the most probable partial sequence at time step *t*, ensuring that it satisfies the mass constraint for each cell. Each cell independently carries out its calculations. Since the computation of each cell relies on all the cells in its upper left column, we employ a lock mechanism to prevent computational race conditions. Starting from the first column, we select the amino acid whose mass aligns with the mass constraint for each row, filling in the respective cell. After completing its computation, the cell releases its dependency lock, allowing the subsequent cell to utilize its results for further computation. Throughout the column, we perform recursive updates in accordance with the PMC rule. Once all cells in the table have completed their calculations, we can retrieve the value in the last cell, which represents the final sequence we seek.

### 2.3 Model configurations and training process

PrimeNovo utilizes an encoder-decoder transformer network configuration. More precisely, both the encoder and decoder components consist of 9 multi-head attention layers. Each attention layer within the network is composed of 8 heads. The hidden embedding dimension for our model is set to 400.

For the training process, we employed a Cosine learning rate scheduler in conjunction with the AdamW optimizer. To mitigate the risk of excessive overfitting, we set the dropout rate at 0.18.

During inference, we use the PMC with precision *e* of 0.1Da. This precision constraint ensures that the decoded peptide always falls within a range of 0.1 Da from the precursor mass indicated in the spectrum.

### 2.4 Peptide annotation for metaproteomic research

The quality assessment of decoded peptides from both PrimeNovo and Casanovo V2, when applied to unlabeled spectra, encompassed various facets, including peptide counts, taxonomic annotations, and functional annotations. Taxonomic annotation was conducted utilizing Unipept (version 5.1.2, employing UniProt 2023.04), while functional annotation was carried out using eggNOG-mapper v2 (utilizing the eggNOG 5 database). To construct the taxon-function network, we employed a protein-peptide bridge approach with the assistance of a pre-existing Python script [15].

The quality control process, denoted as T\U\D\DS, involves the following steps: first, we identify sequences present in the target database. Then, we filter out results that are: (1) unmatched with the precursor mass (mass error larger than 0.1 Da); (2) found within the decoy database; (3) identified in database search results. Both the target and decoy databases were provided in the original study [12]. Additionally, the T\U\D approach is similar but does not entail a comparison with the database search results.

## 3 Additional results on nine-species benchmark dataset

### PrimeNovo extends its strong performance to the revised nine-species benchmark dataset

The widely accepted nine-species benchmark dataset has faced criticism due to its limited amount of testing data. In an effort to address this limitation and provide a more comprehensive evaluation of model performance, Yilmaz et al. [2] introduced a new nine-species test dataset in conjunction with the Casanovo V2 model. This updated test set offers a significantly larger volume of data points compared to its predecessor and features a broader distribution of data across each species. In our study, we conducted tests on this new nine-species dataset using PrimeNovo and compared its performance with that of Casanovo V2. As illustrated in Supplementary Fig. 2, Fig. 3 and Table 5, PrimeNovo consistently outperforms Casanovo V2 across all test species, achieving an average peptide recall rate that surpasses Casanovo V2 by 10%. This outcome reinforces the superior accuracy and efficacy of our model.

### PrimeNovo Achieves State-of-the-Art Amino Acid Level Accuracy

While sequence-level accuracy, measured by peptide recall, remains a paramount metric for evaluating model performance, amino acid (AA) level accuracy holds significance for de novo models. Partially correct amino acids can often provide valuable biological insights and aid in identifying essential components within a peptide. In our comparison of AA level accuracy between PrimeNovo and all other baseline models using the nine-species benchmark dataset, PrimeNovo consistently outperforms its counterparts in AA precision, as illustrated in Supplementary Fig. 4.

Furthermore, we delve into the AA level accuracy at various confidence levels of the model. We rank predictions from both PrimeNovo and Casanovo V2 based on their confidence scores and assess amino acid accuracy within each confidence level range. As demonstrated in Supplementary Fig. 5, amino acid accuracy increases with higher confidence scores for both PrimeNovo and Casanovo. This indicates that model confidence scores serve as meaningful indicators of token-level accuracy. PrimeNovo exhibits superior AA level accuracy in the high-confidence region compared to Casanovo V2, highlighting its ability to make higher-quality token-level predictions when confident in its own predictions. For detailed metrics regarding peptide level accuracy and amino acid level accuracy on the nine-species benchmark dataset, please refer to Supplementary Tables 3 and 4.

**Supplementary Table 3:** Peptide recall on the nine-species benchmark dataset. Note: the bold text indicates the highest performance in each row.

| Species | Peaks. | PepNet | Deep. | Point. | Casa. | Casa.V2 | Prime. |
| --- | --- | --- | --- | --- | --- | --- | --- |
| *A. mellifera* | 0.29 | 0.28 | 0.33 | 0.40 | 0.41 | 0.49 | **0.60** |
| *B. subtilis* | 0.39 | 0.42 | 0.45 | 0.52 | 0.54 | 0.62 | **0.72** |
| *C. endoloripes* | 0.20 | 0.24 | 0.25 | 0.30 | 0.33 | 0.45 | **0.53** |
| *H. sapiens* | 0.28 | 0.23 | 0.29 | 0.35 | 0.34 | 0.45 | **0.57** |
| *M.mazei* | 0.36 | 0.38 | 0.42 | 0.48 | 0.48 | 0.56 | **0.65** |
| *M. musculus* | 0.20 | 0.28 | 0.29 | 0.36 | 0.43 | 0.48 | **0.57** |
| *S. cerevisiae* | 0.43 | 0.37 | 0.46 | 0.53 | 0.49 | 0.60 | **0.70** |
| *S. lycopersicum* | 0.40 | 0.43 | 0.45 | 0.51 | 0.52 | 0.62 | **0.70** |
| *V. mungo* | 0.36 | 0.32 | 0.44 | 0.51 | 0.51 | 0.59 | **0.70** |
| Average | 0.32 | 0.33 | 0.38 | 0.44 | 0.45 | 0.54 | **0.64** |

**Supplementary Fig. 2:**
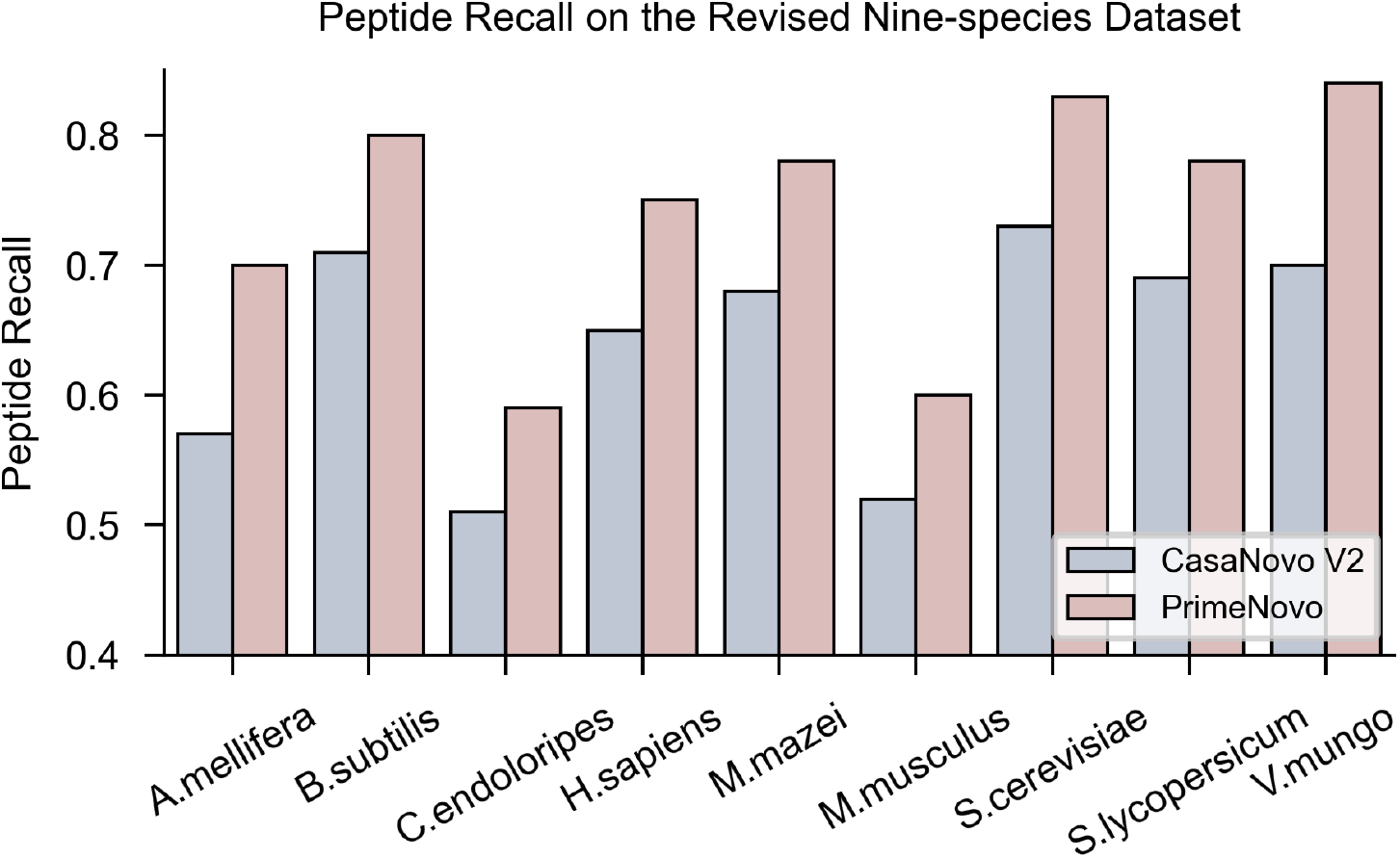
Peptide recall on the revised nine-species dataset.

**Supplementary Fig. 3:**
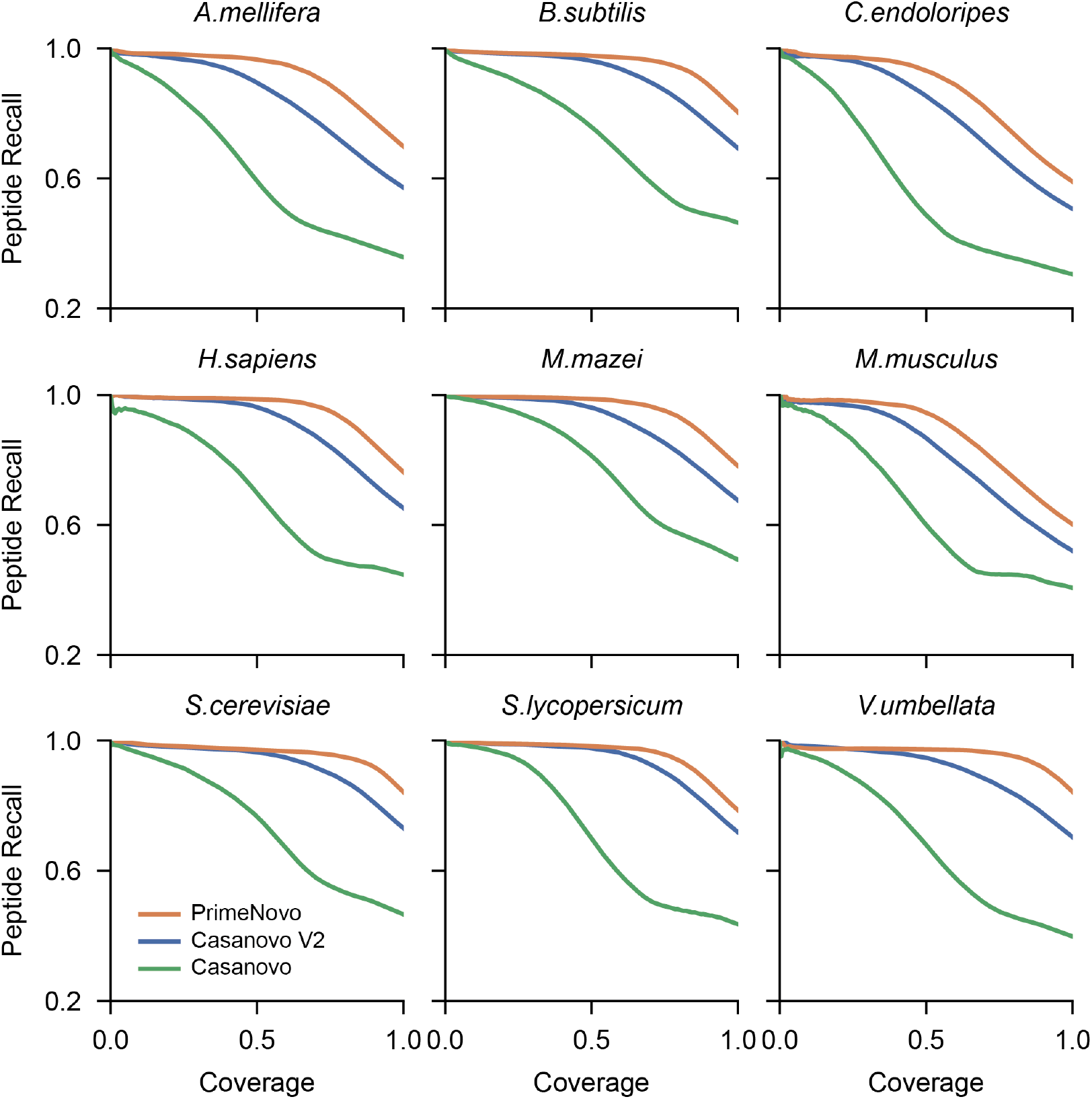
Peptide recall-coverage curves on the revised nine-species dataset.

**Supplementary Fig. 4:**
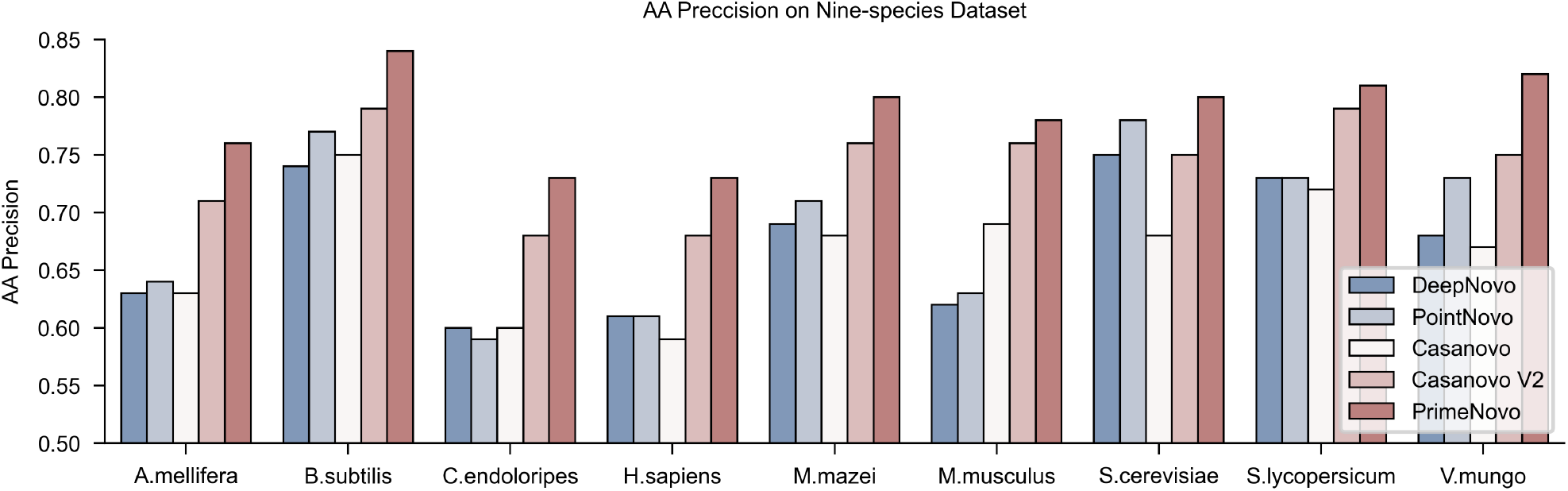
The prediction precision at AA level on the nine-species benchmark dataset. The AA precision of PrimeNovo is significantly higher than that of other de novo methods.

**Supplementary Fig. 5:**
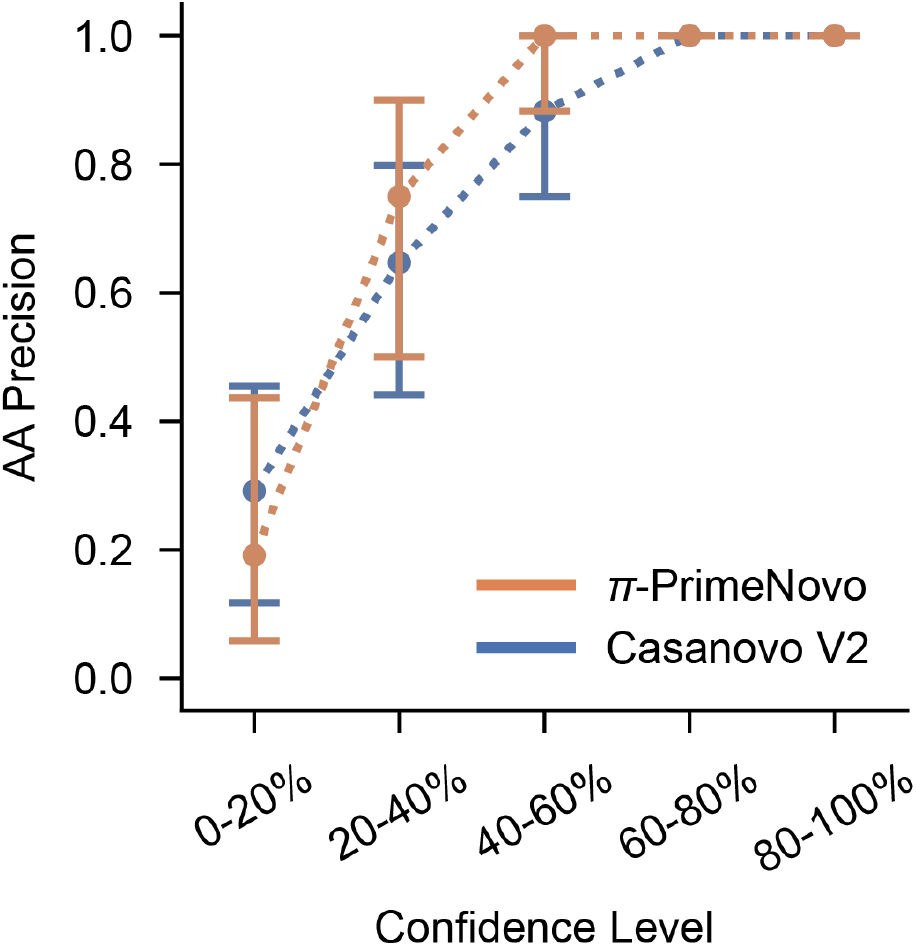
The prediction precision at AA level with the increase of confidence score on nine-species benchmark dataset. PrimeNovo exhibits an overall higher AA precision compared to Casanova V2.

**Supplementary Table 4:**
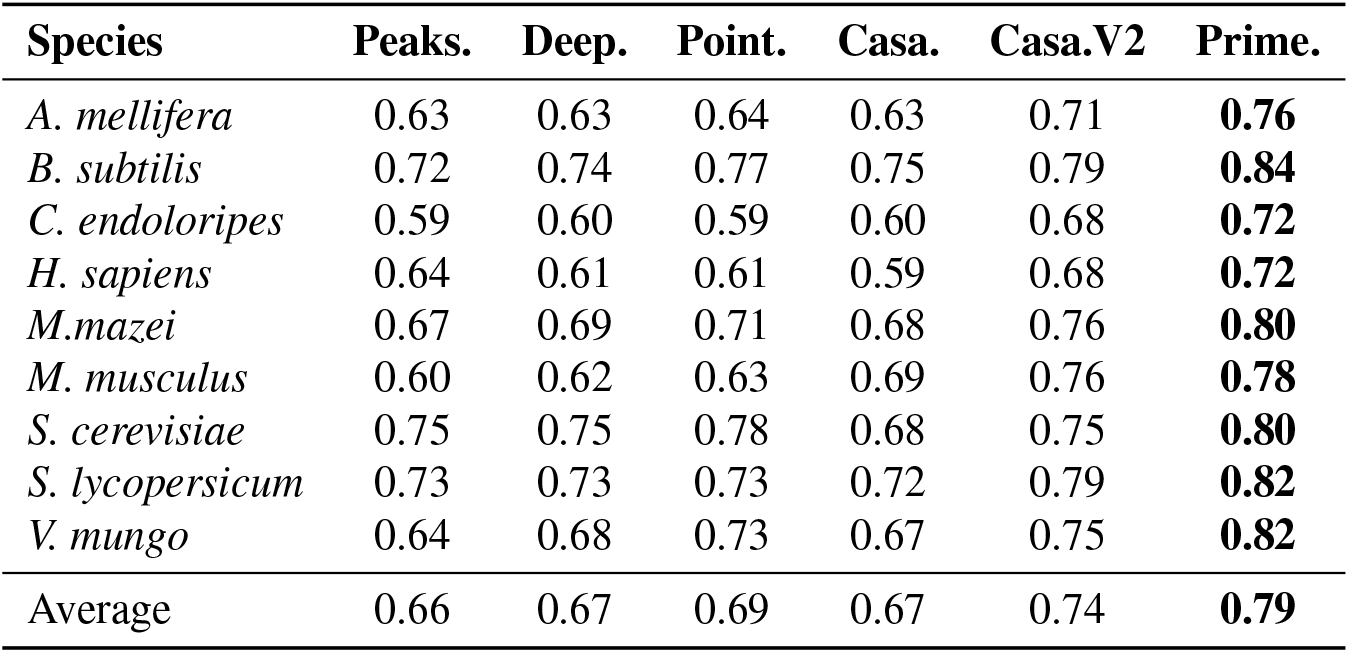
Amino acid precision on the nine-species benchmark dataset. Note: the bold text indicates the highest performance in each row.

| Species | Peaks. | Deep. | Point. | Casa. | Casa.V2 | Prime. |
| --- | --- | --- | --- | --- | --- | --- |
| *A. mellifera* | 0.63 | 0.63 | 0.64 | 0.63 | 0.71 | **0.76** |
| *B. subtilis* | 0.72 | 0.74 | 0.77 | 0.75 | 0.79 | **0.84** |
| *C. endoloripes* | 0.59 | 0.60 | 0.59 | 0.60 | 0.68 | **0.72** |
| *H. sapiens* | 0.64 | 0.61 | 0.61 | 0.59 | 0.68 | **0.72** |
| *M.mazei* | 0.67 | 0.69 | 0.71 | 0.68 | 0.76 | **0.80** |
| *M. musculus* | 0.60 | 0.62 | 0.63 | 0.69 | 0.76 | **0.78** |
| *S. cerevisiae* | 0.75 | 0.75 | 0.78 | 0.68 | 0.75 | **0.80** |
| *S. lycopersicum* | 0.73 | 0.73 | 0.73 | 0.72 | 0.79 | **0.82** |
| *V. mungo* | 0.64 | 0.68 | 0.73 | 0.67 | 0.75 | **0.82** |
| Average | 0.66 | 0.67 | 0.69 | 0.67 | 0.74 | **0.79** |

**Supplementary Table 5:** Peptide recall on the revised nine-species benchmark dataset. Note: the bold text indicates the highest performance in each row.

| Species | Casa. | Casa.V2 | Prime. |
| --- | --- | --- | --- |
| *A. mellifera* | 0.3578 | 0.57 | **0.70** |
| *B. subtilis* | 0.4637 | 0.71 | **0.80** |
| *C. endoloripes* | 0.3055 | 0.51 | **0.59** |
| *H. sapiens* | 0.4468 | 0.65 | **0.75** |
| *M.mazei* | 0.4927 | 0.68 | **0.78** |
| *M. musculus* | 0.4063 | 0.52 | **0.60** |
| *S. cerevisiae* | 0.4657 | 0.73 | **0.83** |
| *S. lycopersicum* | 0.4354 | 0.69 | **0.78** |
| *V. mungo* | 0.3977 | 0.7 | **0.84** |
| Average | 0.4191 | 0.64 | **0.74** |

## 4 Additional results on other test datasets

### PrimeNovo Demonstrates Superior Amino Acid Prediction Across Diverse Unseen Datasets

We also conducted a comparison of PrimeNovo with other de novo algorithms based on amino acid (AA) level precision across three additional datasets. Notably, all models were evaluated under the zero-shot setting, without any fine-tuning on the target data distribution. As depicted in Supplementary Fig. 6a and 6b, along with the results provided in Supplementary Table 6, PrimeNovo excels in AA level accuracy, surpassing the previous best model, in both the HCC and PT datasets. Furthermore, PrimeNovo outperforms all baseline models on the three-species dataset across all tested species, as indicated in Supplementary Table 6. To provide a more detailed analysis, we compared AA level accuracy within each confidence range, having ranked the peptides based on their output confidence scores. Our observations reveal that PrimeNovo consistently achieves higher accuracy levels across all confidence ranges compared to Casanovo V2. These results collectively highlight PrimeNovo’s superior capability in accurately predicting amino acids across a range of unseen datasets, even when operating in a zero-shot setting.

### PrimeNovo Demonstrates Robust Performance in the Face of Missing Peaks and Varied Peptide Lengths

We conducted an analysis of PrimeNovo’s performance, as measured by peptide recall, when dealing with peptides of different lengths and input spectra containing missing data. To accomplish this, we utilized the PT dataset and categorized the MS/MS data based on the length of the target peptide. Within each length category, we quantified the number of missing peaks in each spectrum, following the methodology outlined in previous research [16]. Subsequently, we reported the peptide-level recall under a zero-shot setting for each combination of target peptide length and missing peak count. As observed in Supplementary Fig. 10 and Fig. 8, both models experience a drop in accuracy (indicated by lighter colors) when more peaks are missing from the spectrum, regardless of the peptide length. However, the heat map illustrating the performance gap between PrimeNovo and Casanovo V2 highlights that PrimeNovo consistently maintains a significantly higher level of peptide-level accuracy across all combinations of missing peaks and target lengths. Furthermore, PrimeNovo consistently outperforms Casanovo V2 across all target lengths, underscoring its robustness in handling variations in peptide length as well as the presence of missing peaks in the input data.

### PrimeNovo Demonstrates Enhanced Adaptability on the PT Dataset

In our investigation, we conducted finetuning experiments on both Casanovo V2 and PrimeNovo using the PT dataset. The objective was to evaluate the adaptability of both models to unseen data distributions. Similar to the observations in the “Results” section of our main manuscript, when the model is exclusively fine-tuned using the PT training data, we observe a phenomenon known as catastrophic forgetting, where the model’s performance on the original data distribution significantly declines. This is evident in the left portion of Supplementary Fig. 9, where the addition of more PT data results in a drop in performance on the nine-species benchmark dataset. To address this issue, we implemented a strategy of mixing the training data from PT with the original MassiveKB training data in a proportional manner. As illustrated in the right-hand side of Supplementary Fig. 9, fine-tuning with a mixture of the original training data and PT data led to stable performance on the original nine-species dataset. Moreover, with the gradual addition of more PT training data, performance on the PT test set consistently improved. Crucially, throughout these finetuning experiments with varying dataset sizes and mixing ratios (10:1, 1:1, and 1:10), PrimeNovo consistently outperformed Casanovo V2 across all configurations. This robust performance underlines PrimeNovo’s superior adaptability to different data distributions and highlights its strong generalization capabilities.

### PrimeNovo Excels in Predicting Peptides with Various Charges, Outperforming Recently Published Pep-Net

We conducted a comparison of PrimeNovo’s prediction accuracy with the most advanced CNN-based deep learning model, PepNet [10]. For this analysis, we utilized the PXD019483 dataset, which was employed as the test set by the authors, and organized predictions based on the charges of each peptide. As demonstrated in Supplementary Fig. 11a, PrimeNovo attains the highest peptide-level accuracy across peptides with different charges, outperforming PepNet. Additionally, we compared the performance of the PepNet model with PrimeNovo on the nine-species benchmark dataset, as depicted in Supplementary Fig. 11b. In this evaluation as well, PrimeNovo exhibits a significant advantage in terms of peptide-level recall over PepNet, underscoring its superior predictive capabilities.

**Supplementary Fig. 6:**
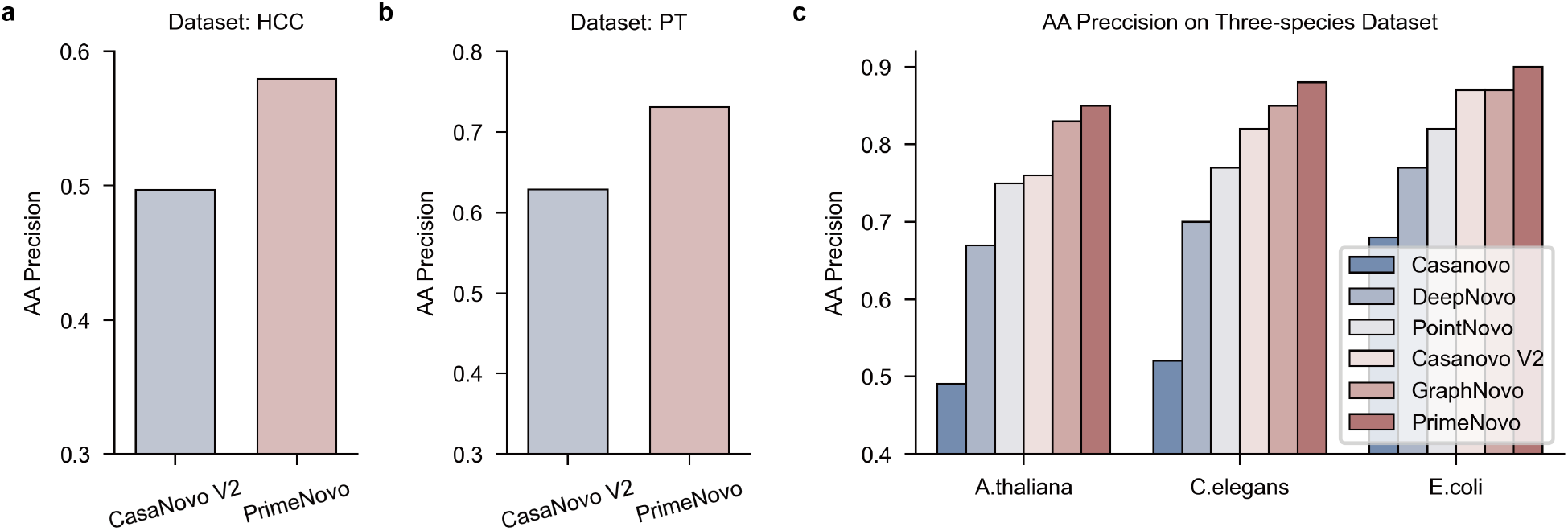
The prediction precision at Amino Acid level on the test datasets HCC (a), PT (b), and Three-species (c), respectively.

**Supplementary Fig. 7:**
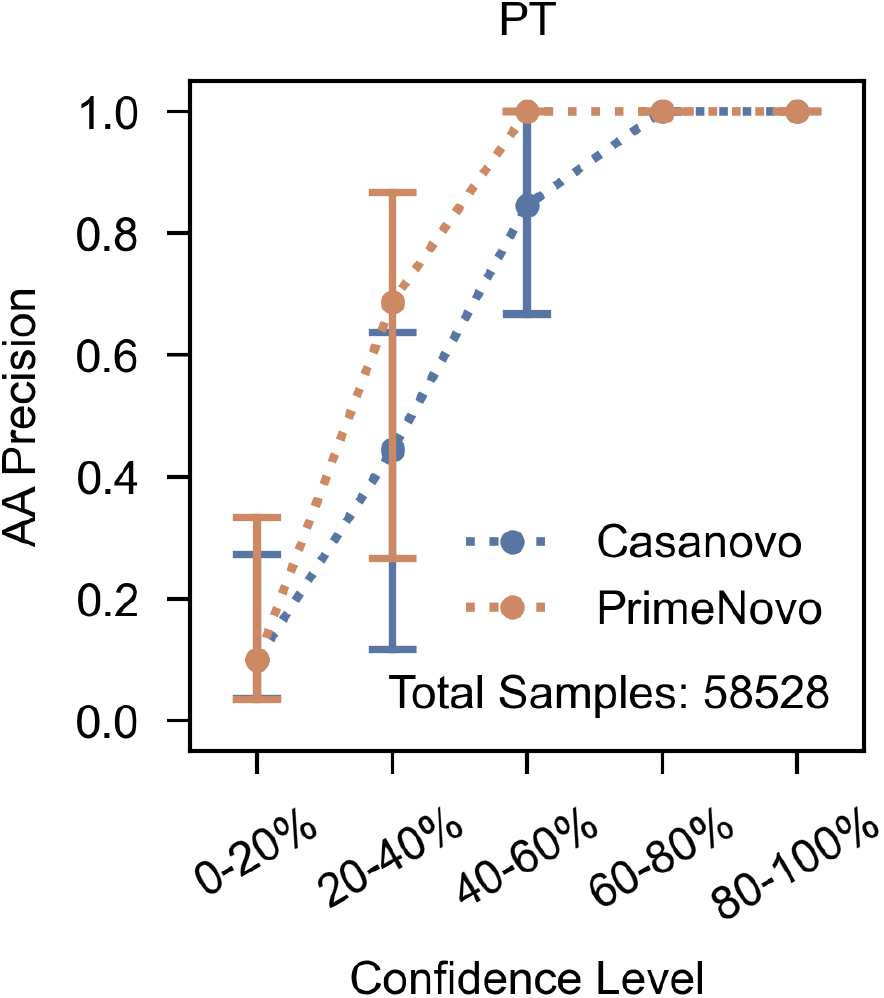
The prediction precision at AA level with the increase of confidence score on the PT dataset. PrimeNovo exhibits an overall higher AA precision compared to Casanova V2.

**Supplementary Table 6:** The average performance of PrimeNovo compared to Casanovo and Casanovo V2 across four distinct large-scale MS/MS datasets.

| Test dataset | Amino acid precision |  |  | Peptide recall |  |  |
| --- | --- | --- | --- | --- | --- | --- |
|  | Casanovo | Casanovo V2 | PrimeNovo | Casanovo | Casanovo V2 | PrimeNovo |
| HCC | 0.0908 | 0.4968 | 0.5793 | 0.0001 | 0.161 | 0.3817 |
| IgG1-Human-HC | 0.34465 | 0.605916667 | 0.693133333 | 0.1154 | 0.4065 | 0.5416 |
| PT | 0.4443 | 0.6284 | 0.7307 | 0.2873 | 0.4543 | 0.5857 |
| Three-species | 0.5633 | 0.8167 | 0.8767 | 0.5037 | 0.6873 | 0.7867 |

**Supplementary Fig. 8:**
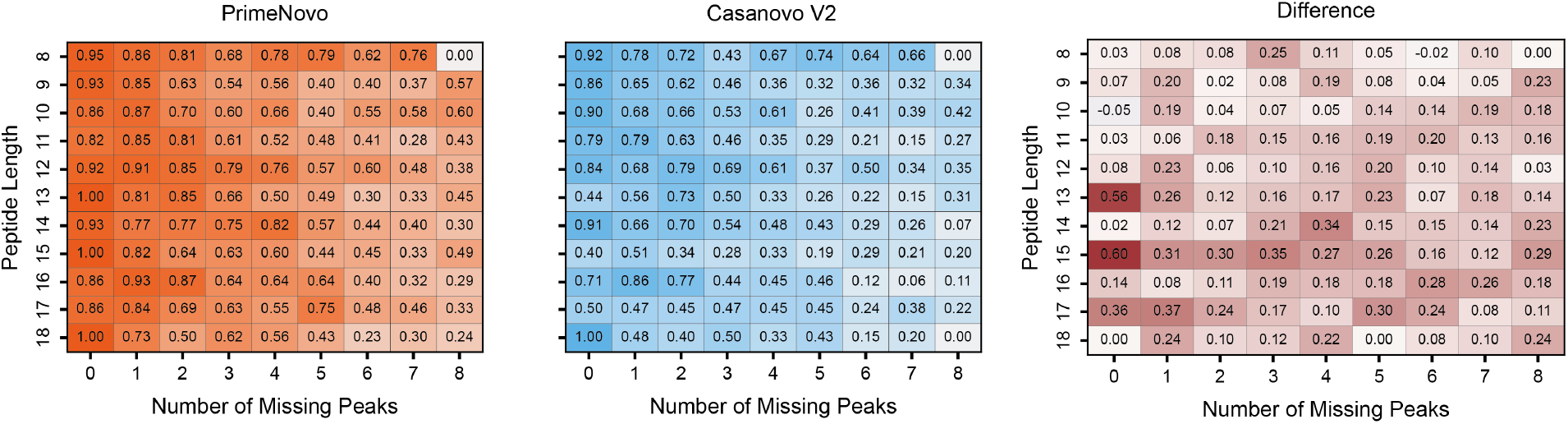
The peptide recalls are shown by heatmap for different combinations of the number of missing peaks and peptide lengths of PrimeNovo (left) and Casanovo V2 (middle). The right heatmap shows the differences in peptide recalls between PrimeNovo and Casanovo V2 for different combinations of the number of missing peaks and peptide lengths.

**Supplementary Fig. 9:**
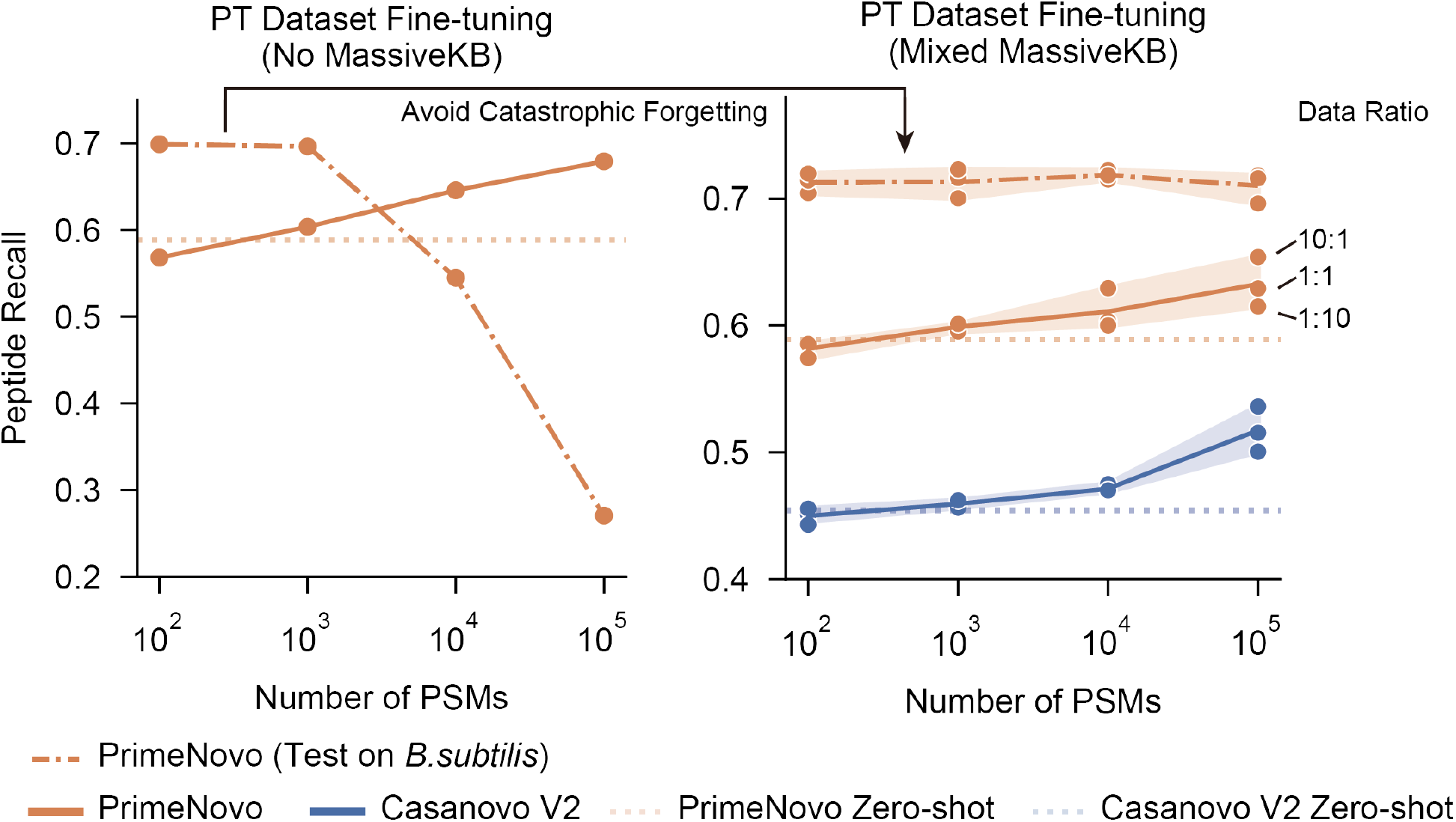
The performance of fine-tuning on the PT test dataset as more PT training data is added during fine-tuning. The left panel shows fine-tuning using only the PT dataset, which leads to a catastrophic forgetting of the original data distribution (on part of the nine-species benchmark dataset *B*.*subtilis*). The right side illustrates fine-tuning with a mixture of PT and MassIVE-KB training data.

**Supplementary Table 7:** The performance of PrimeNovo compared to Casanovo and Casanovo V2 on six different proteolytic enzymes in the IgG1-Human-HC dataset.

| Enzyme | Amino acid precision |  |  | Peptide recall |  |  |
| --- | --- | --- | --- | --- | --- | --- |
|  | Casanovo | Casanovo V2 | PrimeNovo | Casanovo | Casanovo V2 | PrimeNovo |
| AspN | 0.3089 | 0.5236 | 0.6059 | 0.0726 | 0.2611 | 0.3593 |
| Chymotrysin | 0.2114 | 0.4794 | 0.6227 | 0.0298 | 0.2493 | 0.446 |
| GluC | 0.2878 | 0.5214 | 0.6427 | 0.0686 | 0.3133 | 0.4625 |
| LysC | 0.4359 | 0.7002 | 0.749 | 0.2108 | 0.5224 | 0.6204 |
| ProteinaseK | 0.3844 | 0.6735 | 0.7567 | 0.089 | 0.4568 | 0.5972 |
| Trypsin | 0.4395 | 0.7374 | 0.7818 | 0.1703 | 0.5483 | 0.6649 |

**Supplementary Fig. 10:**
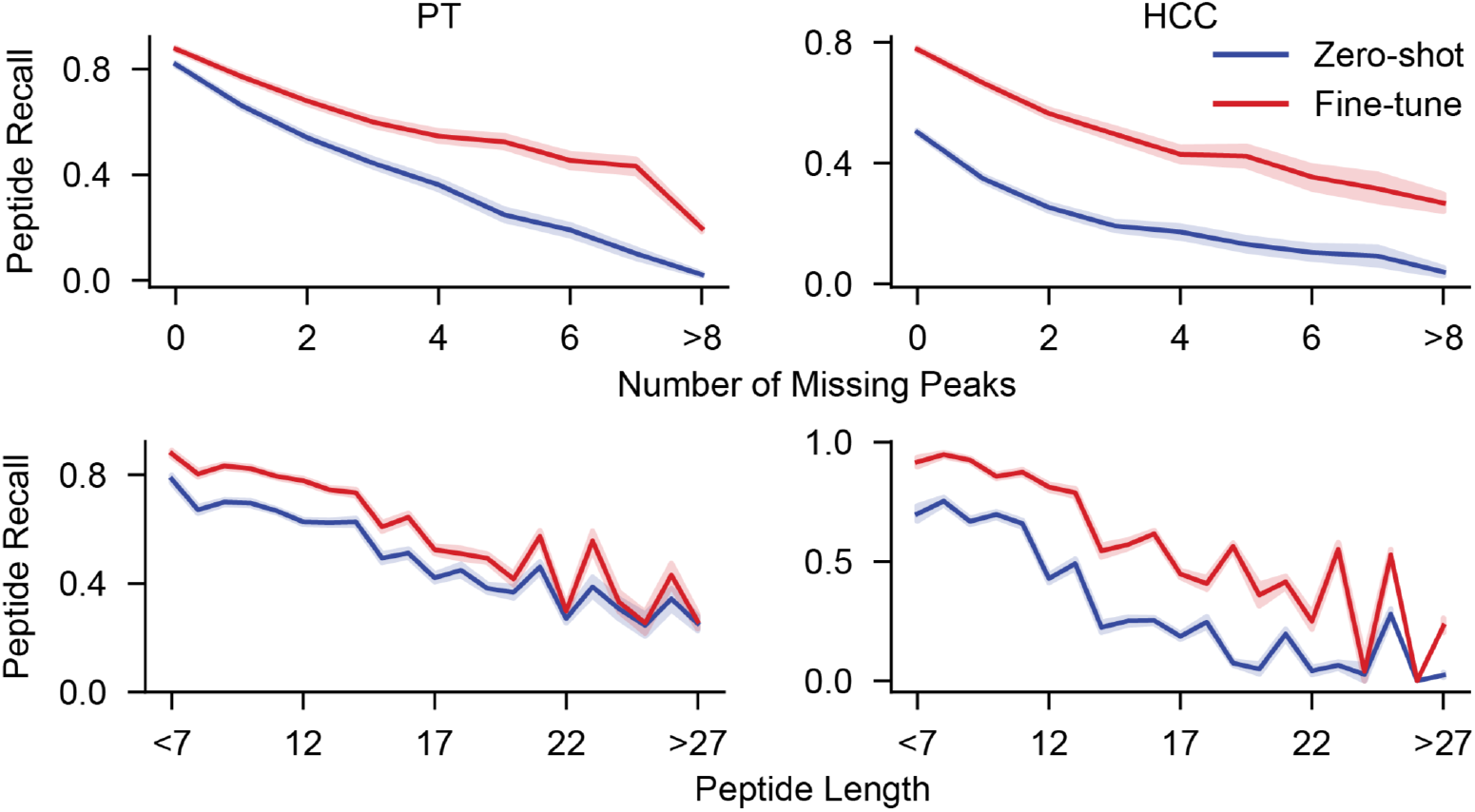
The performance of PrimeNovo for varying peptide lengths and missing peaks.

**Supplementary Fig. 11:**
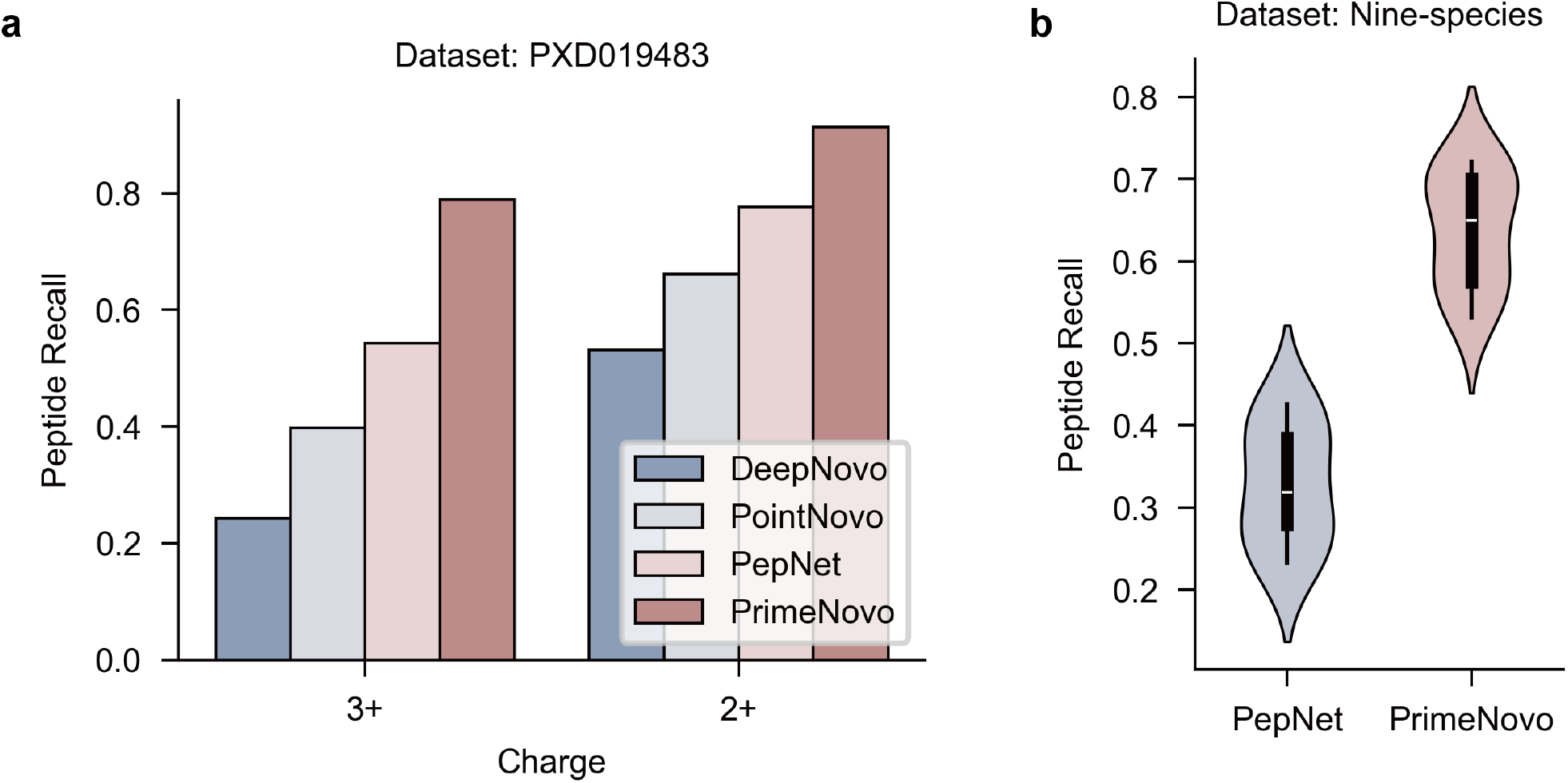
Performance comparison with PepNet. **a**. Comparing PrimeNovo with DeepNovo, PointNovo and PepNet on the PepNet(Human) dataset at 2+ and 3+ charges, respectively. **b**. Comparison of PrimeNovo on the nine-species benchmark dataset with PepNet.

**Supplementary Table 8:**
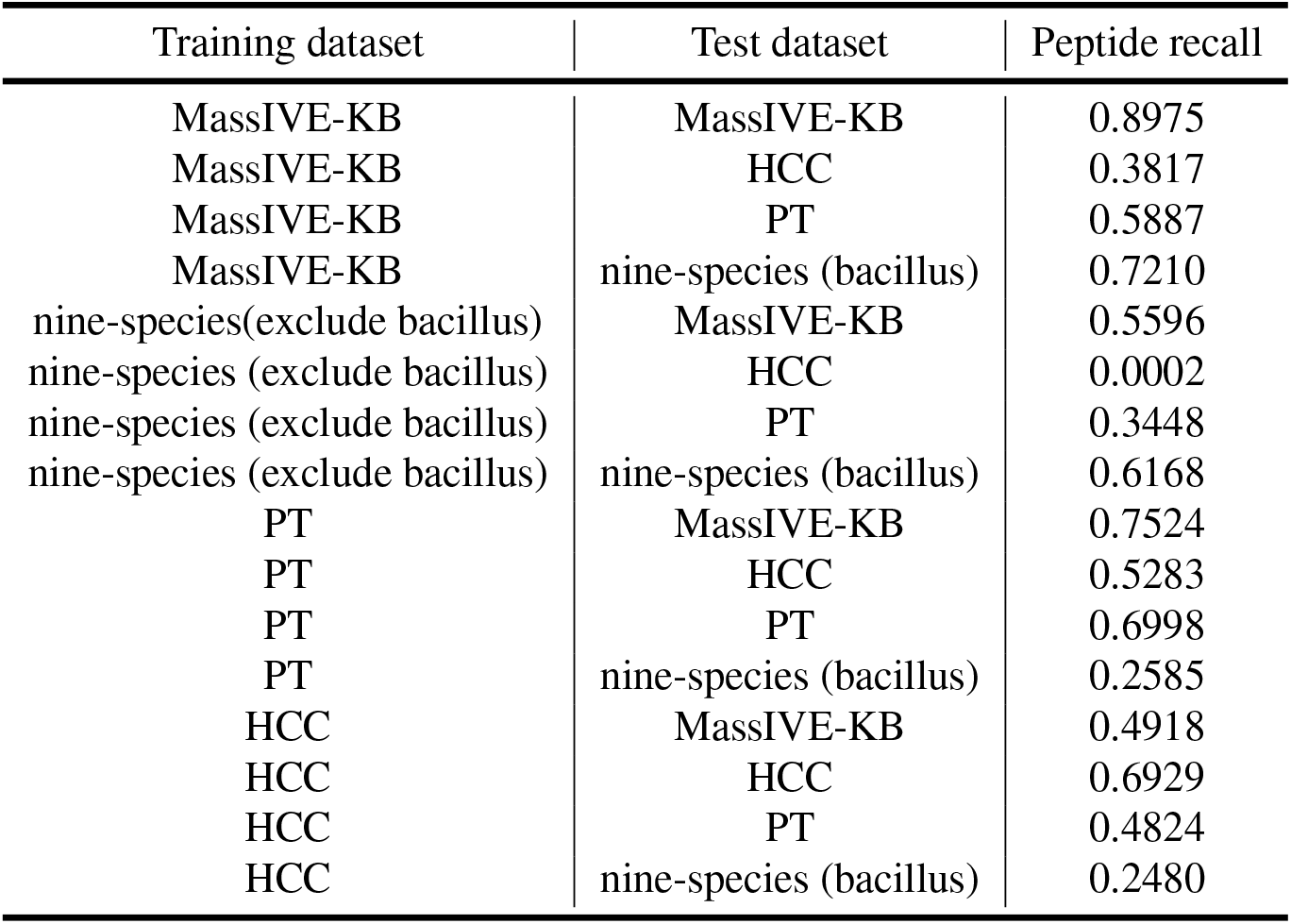
The average peptide recalls when assessing the model’s generalization ability.

**Supplementary Table 9:** The performance of PrimeNovo on the phosphorylation dataset 2020-Cell-LUAD.

| Data type | Classification accuracy | AA recall | Peptide recall |
| --- | --- | --- | --- |
| Tumor tissue | 0.98 | 0.80 | 0.66 |
| Non-cancerous adjacent tissue | 0.98 | 0.85 | 0.66 |

**Supplementary Table 10:**
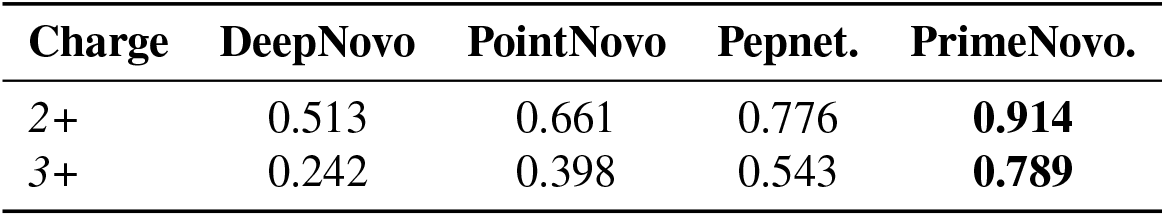
Peptide recall on the Pepnet test dataset under zero-shot setting. The performance for the different precursor charges is reported separately.

| Charge | DeepNovo | PointNovo | Pepnet. | PrimeNovo. |
| --- | --- | --- | --- | --- |
| 2+ | 0.513 | 0.661 | 0.776 | **0.914** |
| 3+ | 0.242 | 0.398 | 0.543 | **0.789** |

## 5 Additional results on model explanalibity

**Supplementary Fig. 12:**
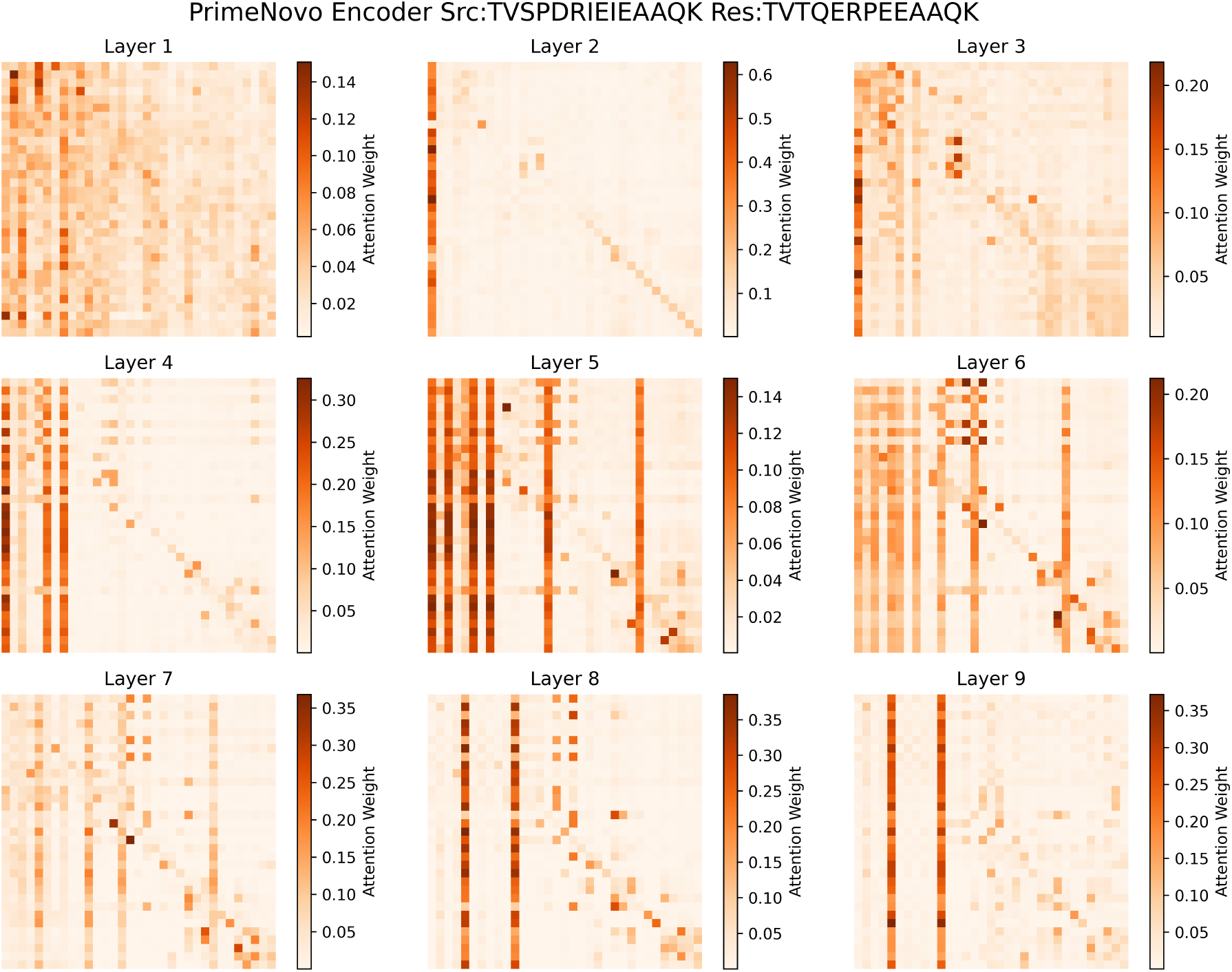
Visualization of the attention score matrix of PrimeNovo’s encoder. A striped distribution in the highlighted sections is revealed. The columns that are highlighted represent the ions selectively preferred by the attention mechanism.

**Supplementary Fig. 13:**
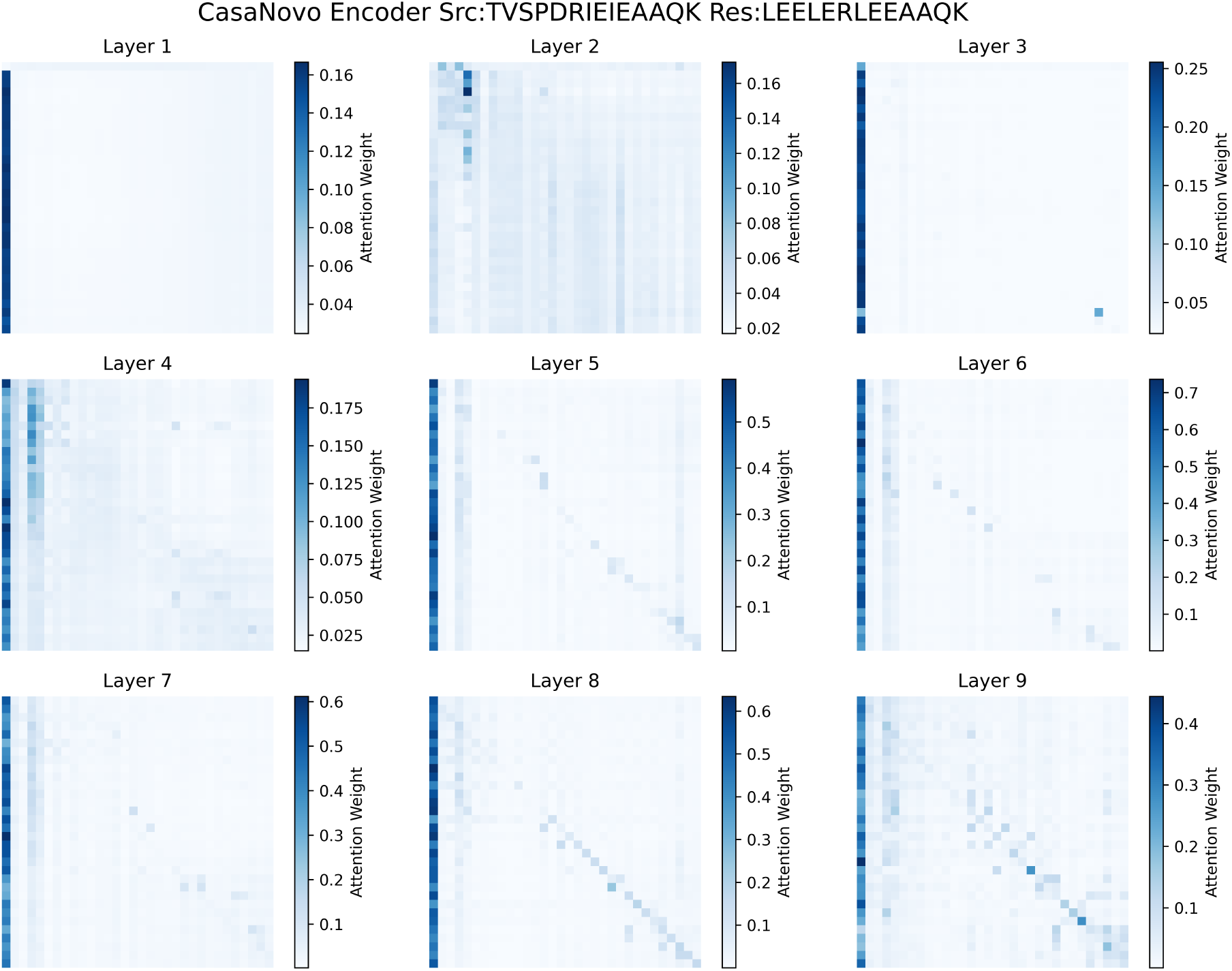
The visualization of the attention score matrix of Casanovo’s encoder. There is no obvious striped distribution in the highlighted sections is revealed. The attention mechanism’s bias for selecting specific ions is not pronounced.

**Supplementary Fig. 14:**
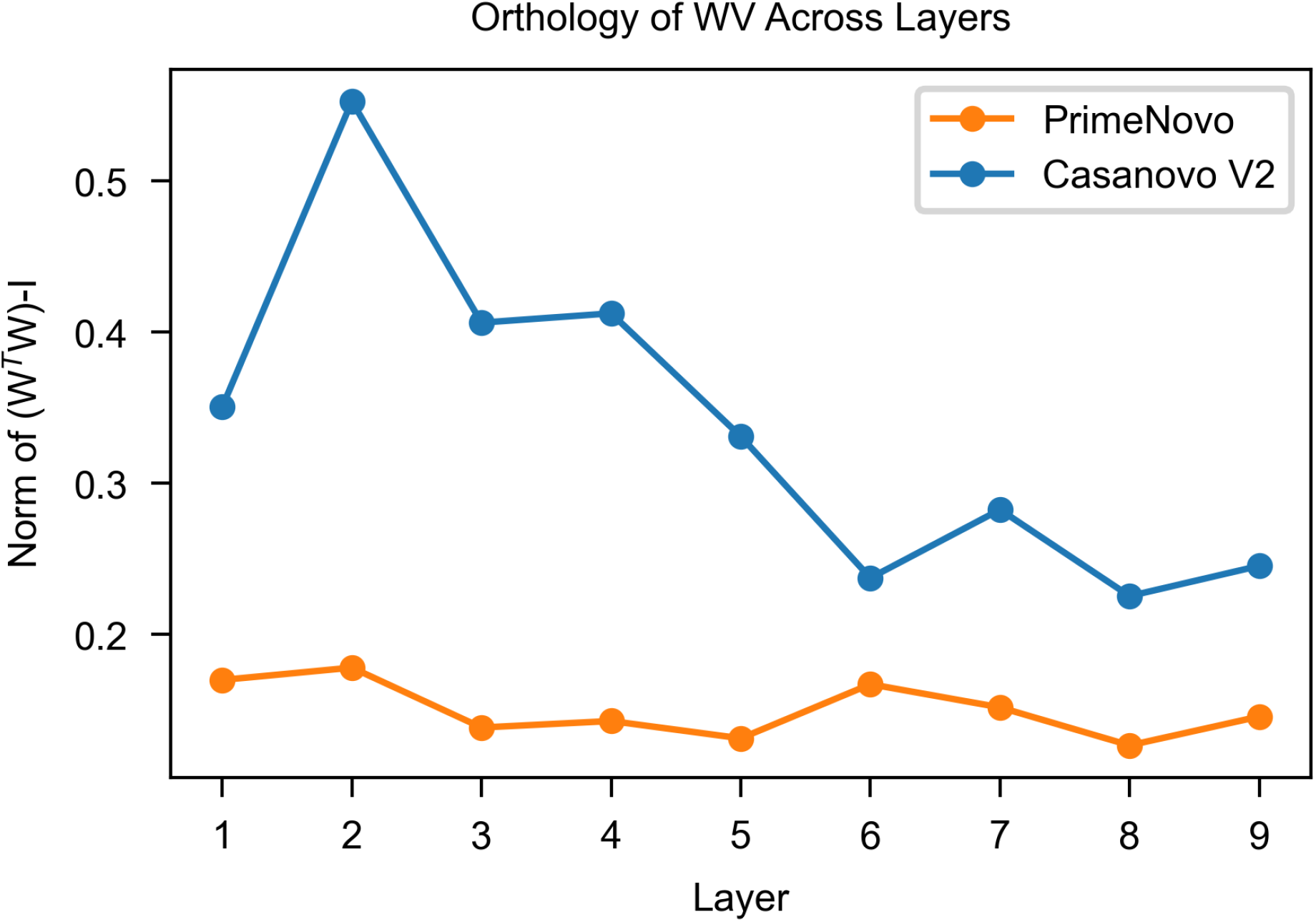
Orthogonality analysis of value matrix projections by the Gram matrix. The y-axis denotes the Frobenius norm of the Gram matrix of the normalized value matrix after subtracting an identity matrix. A smaller value of this norm indicates a better orthogonality of the value matrix projections.

**Supplementary Fig. 15:**
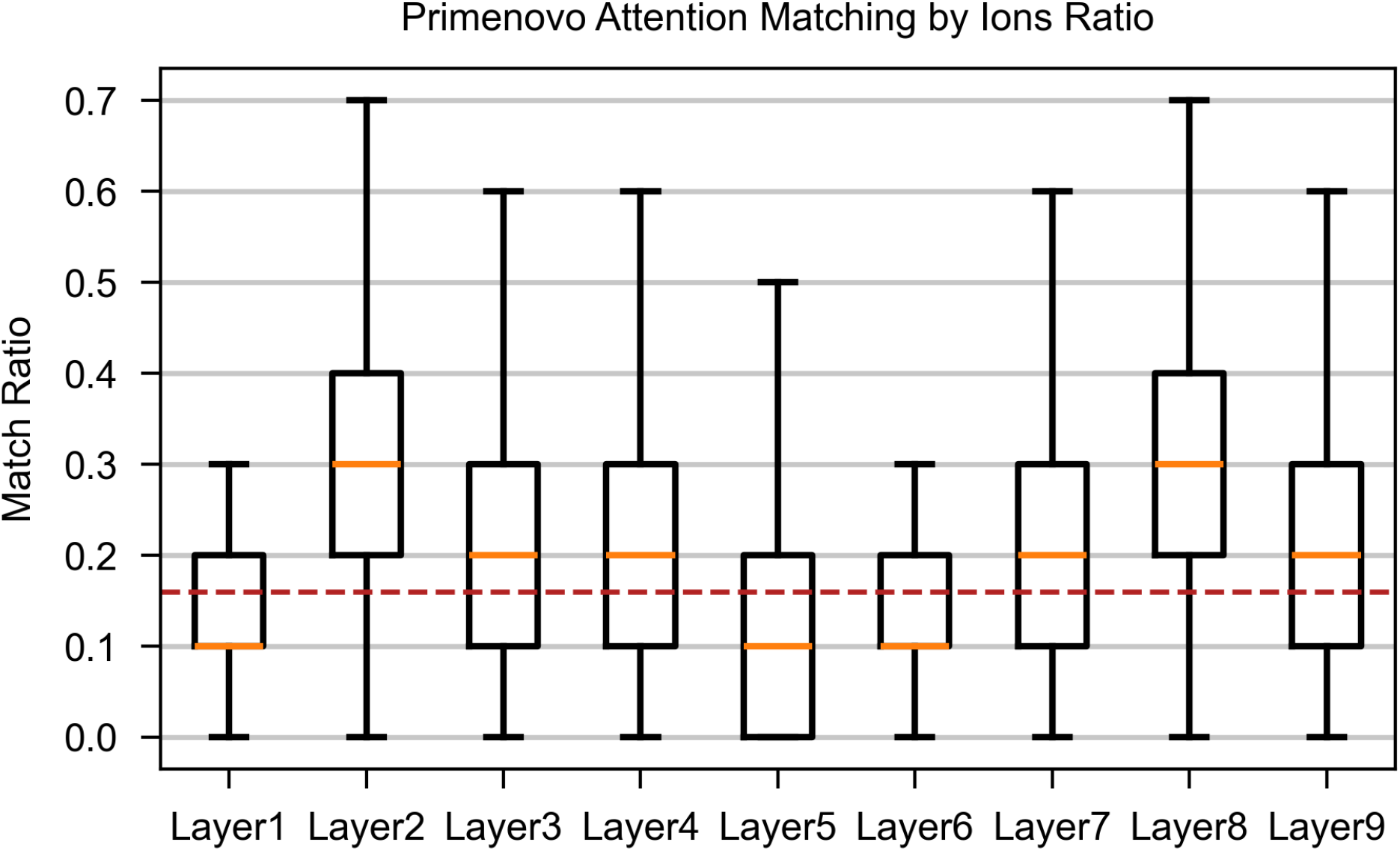
Boxplots to visualize the distribution of the match ratio in the attention score matrices of PrimeNovo’s encoder. The yellow line denotes the median, and the upper and lower edges of the box indicate the first and third quartiles, respectively. The short lines at the top and bottom represent the maximum and minimum values. The red dashed line is the proportion of by ions among all peaks.

### Analyzing the orthogonality of the Value matrix projection by the norm of the Gram matrix

In order to maintain consistency with previous studies, we utilized the norm of the Gram matrix to measure the orthogonality of the Value matrix projection. In this methodology, the Value matrices from all heads at each layer are concatenated to form a complete Value matrix. Subsequently, each column of the Value matrix is normalized according to the L2 norm to transform it into a vector with a magnitude of 1. This normalized matrix is then transposed and multiplied by itself to generate a Gram matrix. The measure of orthogonality is quantified by taking the Frobenius norm of the resulting matrix after the subtraction of an identity matrix. A smaller value of this norm indicates better orthogonality of the Value matrix projection, implying reduced redundancy in the extracted features. As shown in Supplementary Fig. 14, we employ a line graph to visualize and compare the orthogonality of the Value matrix projections in the Casanovo and PrimeNovo encoders. Notably, the curve for PrimeNovo consistently lies below that of Casanovo, demonstrating superior orthogonality in the Value matrix projection of the PrimeNovo encoder, which suggests more diverse feature extraction.

### Analyzing the bias of the PrimeNovo encoder’s attention mechanism towards *b* − *y* ions

On the nine-species benchmark dataset, we random select a spectrum as an example and examine the distribution of match ratio for different layers in the PrimeNovo encoder (Supplementary Fig. 12 and 13). Here, match ratio refers to the proportion of by ions in the top 10 peaks (excluding the first token) that receive the most attention in each sample. This attention level is measured by the L1 norm of the corresponding column in the attention score matrix, with a larger norm indicating a higher attention level.

### Visualization of the distribution of match ratio

As shown in Supplementary Fig. 15, we conducted the analysis on the test dataset of H.sapiens, in which the proportion of *b* ions and *y* ions among all peaks is 0.16. This threshold signifies the expected match ratio if the attention mechanism’s bias towards *b* ions and *y* ions were random. A value exceeding this threshold indicates a preference for ions. The majority of layers exhibit a preference for focusing on *b* ions and *y* ions, particularly the second and eighth layers, aligning with the prior knowledge that ions contain crucial information for de novo sequencing. Conversely, layers 1, 5, and 6 tend to overlook *b* ions and *y* ions, possibly due to their function in extracting useful information from other peaks. A common feature across all layers is the deviation of match ratio from that of random selection, suggesting a definitive selective bias of the attention mechanism towards *b* ions and *y* ions.

### Generation of all possible fragment ions

In this analysis, we considered only the fragment ions generated by the cleavage of chemical bonds along the peptide backbone. It is well-known that in mass spectrometry analysis of peptides, the focus is primarily on six ions generated by the cleavage of three types of chemical bonds. These include a-ions and x-ions formed by the cleavage between the *α*-carbon atom and the carboxyl group, b ions and y ions formed by the cleavage between the carbonyl group and the amino group of the peptide bond, and c ions and z ions formed by the cleavage between the *α*-carbon atom and the amino group. For this study, we also focused on these three types of chemical bond cleavages. However, we not only considered these six ions but also took into account all possible intermediate fragment ions of variable lengths. This process was implemented by the programming language Python.

## 6 Additional results on PTM

**Supplementary Fig. 16:**
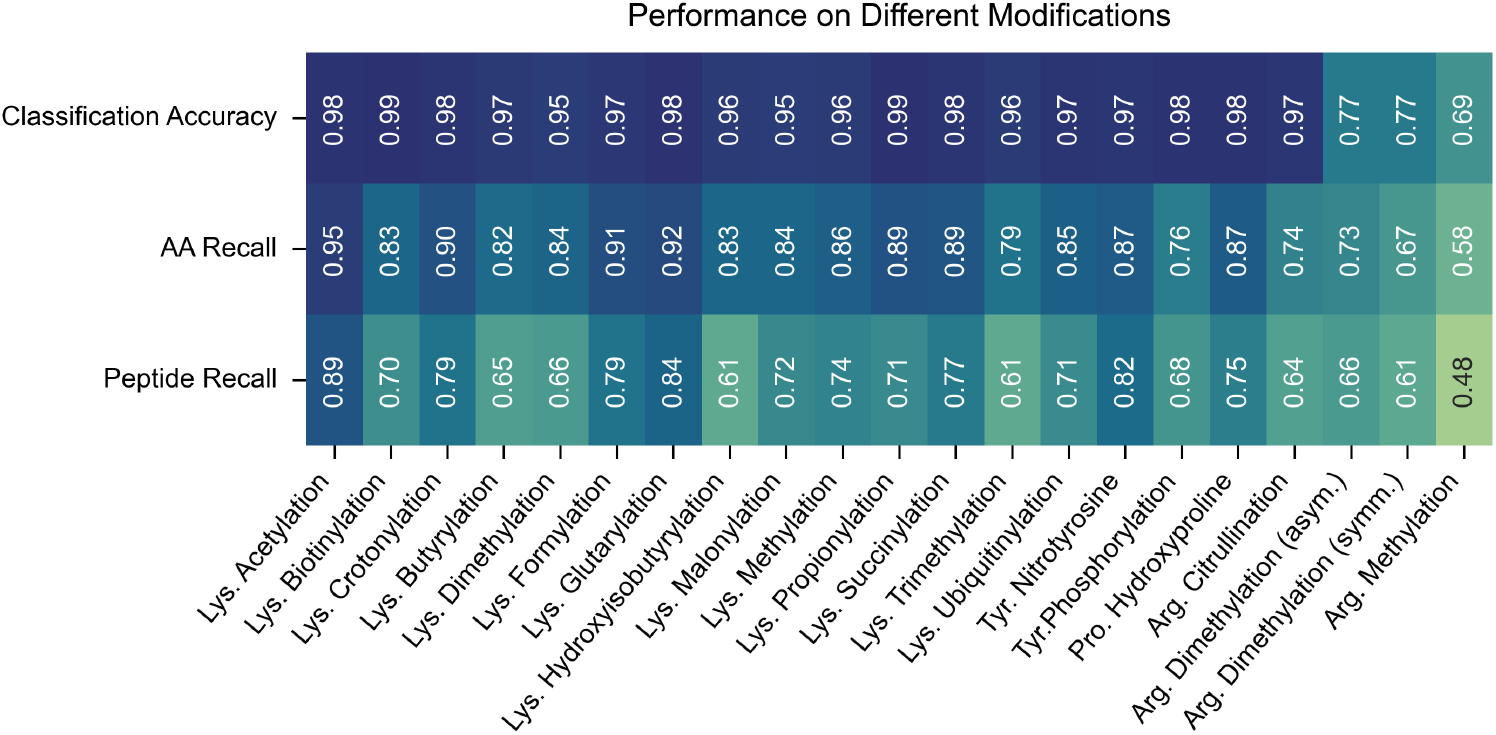
The performance of PrimeNovo for each specific PTM on the 21PTMs dataset.

**Supplementary Fig. 17:**
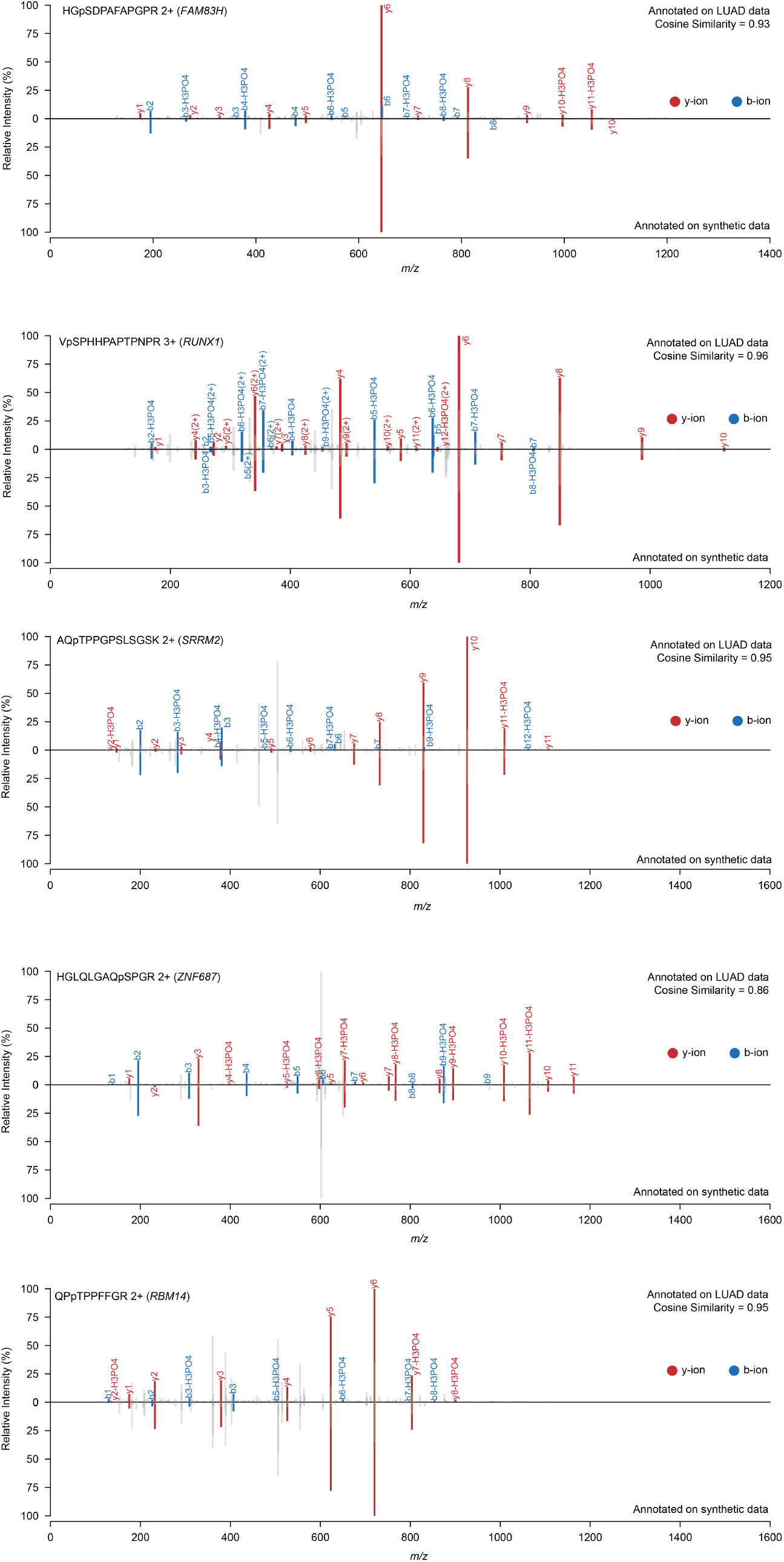
Comparison of the actual input spectrum and spectrum of 10 synthesized peptides predicted by PrimeNovo. The upper section of each diagram displays the original input spectrum from non-enriched 2020-Cell-LUAD dataset, while the lower section showcases the spectrum from the synthetic data. All overlapping peaks are marked in red and blue for b-y ions.

**Supplementary Fig. 18:**
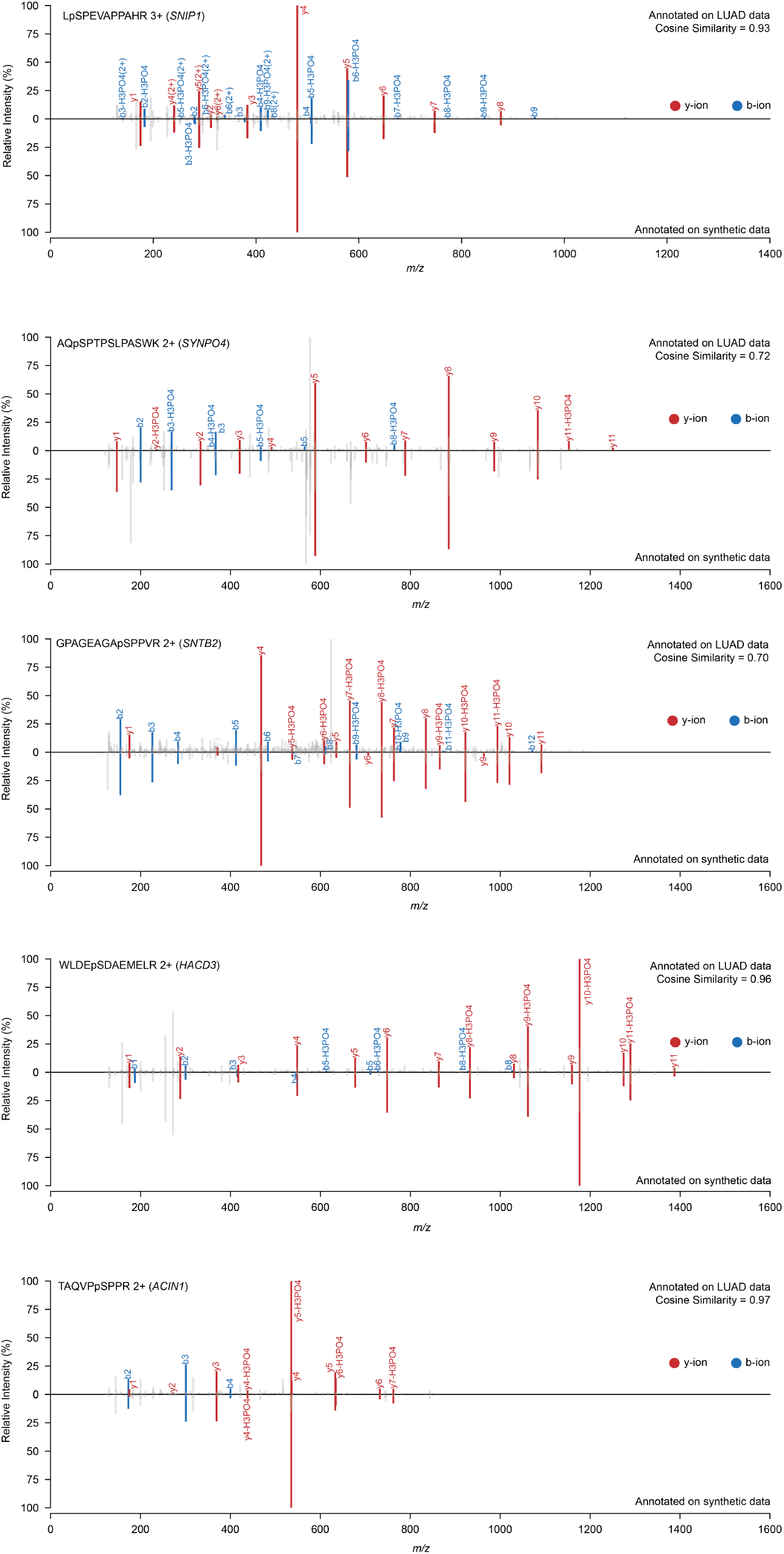
(Cont.) Comparison of the actual input spectrum and spectrum of 10 synthesized peptides predicted by PrimeNovo. The upper section of each diagram displays the original input spectrum from non-enriched 2020-Cell-LUAD dataset, while the lower section showcases the spectrum from the synthetic data. All overlapping peaks are marked in red and blue for b-y ions.

## 7 Additional Result on Synthesizing phospho-peptide from PrimeNovo

### 7.1 Peptide Selection Process

We applied the fine-tuned PrimeNovo model to the non-enriched 2020-Cell-LUAD dataset, notably a dataset without pre-identified peptide labels. The model’s confidence score served as the primary criterion for selection due to the absence of these labels. We initially filtered for the top 300 predicted peptides featuring phospho modifications, all of which boasted confidence scores exceeding 0.99. Subsequent manual scrutiny allowed us to refine this selection based on quality indicators, specifically: 1) Preference for peptides terminating with the amino acids K or R; 2) Selection of peptides of an optimal length, avoiding those excessively long or short; 3) Requirement that more than half of the 20 most intense peaks could be aligned with the theoretical ions of the peptide in question. 4) The presence of a matched ion both preceding and following the phospho modification site.

Following these criteria, we identified 12 peptides as prime candidates for laboratory synthesis and subsequent functional analyses.

### 7.2 Mass Spectrometry Analysis of Synthetic Phosphopeptides

Liquid chromatography tandem mass spectrometry (LC-MS/MS) analyses were conducted using an Orbitrap Fusion Tribrid Lumos mass spectrometer (Thermo Fisher Scientific), directly interfaced with a nanoflow LC system (EASY-nLC 1200, Thermo Fisher Scientific). A quantity of 50 ng of synthetic phosphopeptides was resuspended in solvent A (0.1% formic acid (FA) in HPLC-grade water) and introduced onto a 2 cm self-packed trapping column (100-*μ*m inner diameter, 1.9 *μ*m resin, ReproSil-Pur C18-AQ, Dr. Maisch GmbH) in solvent A. Following loading and washing steps, peptides were eluted to a 30 cm analytical column (150-*μ*m inner diameter, 1.9 *μ*m resin, ReproSil-Pur C18-AQ, Dr. Maisch GmbH) and separated using a non-linear gradient of solvent B (0.1% FA in acetonitrile, ACN) over 30 minutes at a flow rate of 600 nL/min. The gradient program was as follows: from 0 to 2 minutes, 7–12% solvent B; 2 to 13 minutes, 12–32% solvent B; 13 to 19 minutes, 32–45% solvent B; 19 to 21 minutes, 45–95% solvent B; and maintaining 95% solvent B from 21 to 30 minutes. Ionization was achieved with a 2.0 kV spray voltage and a capillary temperature set at 320 °C. Data acquisition was carried out in OT-OT mode with full MS scans (350 to 1,500 m/z) at a resolution of 120,000, a maximum injection time of 50 ms, and an Automatic Gain Control (AGC) target of 1e6. MS2 fragmentation was performed using higher-energy collision dissociation (HCD) with a normalized collision energy of 32%, on 2+ to 7+ precursor ions within a 3 s duty cycle, in top-speed mode. MS2 spectra were acquired in the ion trap in rapid mode, with an AGC target of 50,000 and a maximum injection time of 22 ms, at a resolution of 15,000. Dynamic exclusion was applied for 25 s to prevent the redundant selection of peptides.

### 7.3 Analysis of MS Data for Synthetic Phosphopeptides

The acquired Thermo raw files were analyzed using MaxQuant (version 2.1.4), querying against the human UniProt database (SwissProt, 20,266 entries, release date 2021-01-22). The analysis incorporated variable modifications such as oxidation on methionine, phosphorylation on serine, threonine, and tyrosine, and acetylation at the N-termini of proteins. Fixed modifications included carbamidomethylation of cysteine. The search parameters permitted up to two missed cleavages by trypsin (with full specificity). To ensure data quality, the Percolator algorithm was employed to maintain the false discovery rate (FDR) below 1% at both the peptide spectrum match (PSM) and protein levels.

### 7.4 Details of 12 synthesized Phosphopeptides

We analyzed each one of 12 phosphopeptides in detail from various aspects, with summarized information in Supplementary Table 11 and Table 12.

**Supplementary Table 11:**
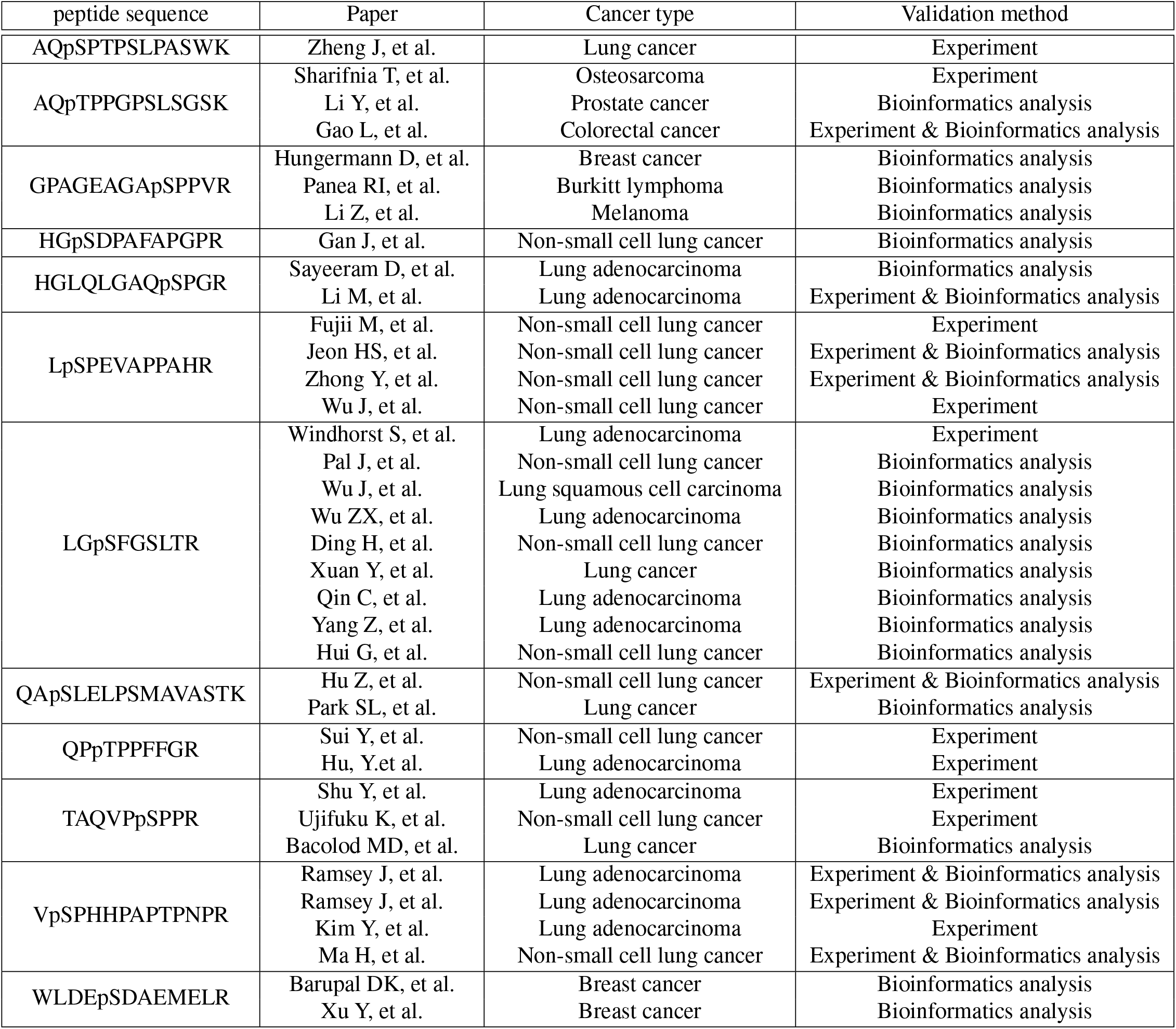
Detailed information about cancer research for the proteins of the 12 synthesized phosphopeptides.

**Supplementary Table 12:**
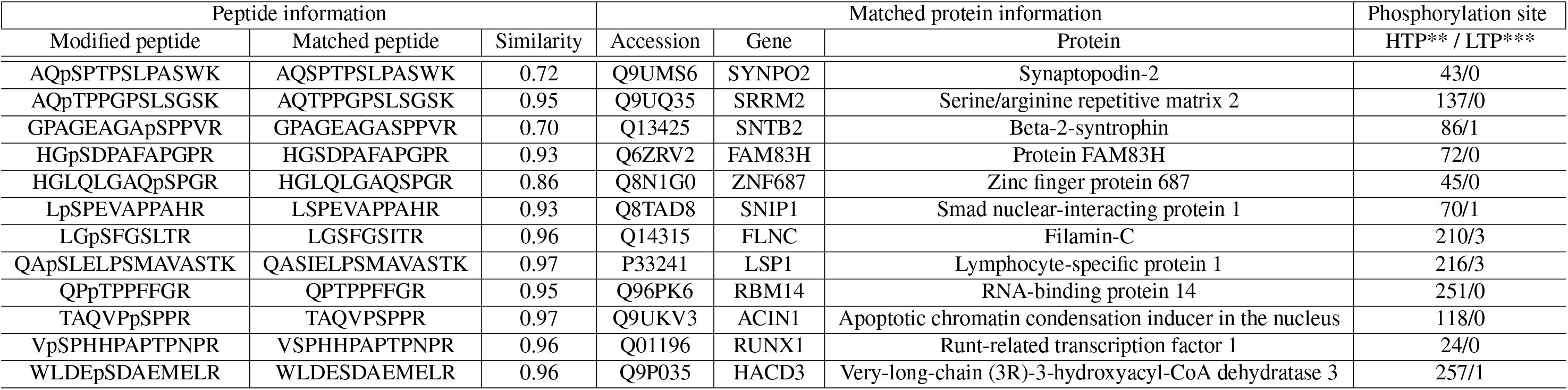
Detailed annotations for all the 12 synthesized phosphopeptides.

### 7.5 More about Three LUAD-Unrelated Peptides from Twelve Selected Predictions

SRRM2 is indispensable for pre-mRNA splicing, serving as a vital component of the spliceosome. Within the minor spliceosome, it potentially contributes to the splicing of U12-type introns found in pre-mRNAs. Research highlights a notable correlation between SRRM2 expression and prostate cancer prognosis [15]. Additionally, SRRM2 possesses phosphorylation sites uniquely regulated by GSK3*α*, significantly impacting the survival outcomes of colorectal cancer patients [17].

SNTB2 is crucial for the subcellular localization and organization of various membrane proteins. It may connect different receptors to the actin cytoskeleton and the dystrophin-glycoprotein complex, influencing the regulation of secretory granules through interactions with PTPRN. Recent studies have identified SNTB2 as a critical immune marker in macrophages, linked to melanoma metastasis [18]. Mutations within the SNTB2 gene are prevalent across Burkitt lymphoma (BL) subtypes, underscoring its potential significance in the disease’s pathology [19].

HACD3 plays a crucial role in the third of four reactions in the long-chain fatty acid elongation cycle. It is involved in producing very long-chain fatty acids (VLCFAs) of various lengths, serving as precursors for membrane lipids and lipid mediators, essential for numerous biological functions. Studies have linked HACD3 to the aggressiveness and prognosis of breast cancer, including its association with patient survival rates [20, 21].

## Footnotes

1 Please refer to the Datasets section in the Supplementary Information for detailed dataset information.

2 Note that we are using Casanovo V2 without Beam Search (BS) due to the estimated inference time with BS exceeding 4000 A100 GPU hours on this large scale dataset, which amounts to more than 21 days of inference with 8 A100 GPUs.

## Notes

### Competing Interest Statement

The authors have declared no competing interest.

### Summary of Updates

revise the email address in the paper for the corresponding author.

